# *Tle4* controls both developmental acquisition and postnatal maintenance of corticothalamic projection neuron identity

**DOI:** 10.1101/2022.05.09.491192

**Authors:** Maria J. Galazo, David Sweetser, Jeffrey D. Macklis

## Abstract

Identities, circuitry, and function of distinct neuron subtypes are specified during embryonic development, then maintained during postnatal maturation and potential plasticity. Mechanisms that control early acquisition of neuron subtype identities, encompassing circuitry and function, in the cerebral cortex have become increasingly understood. However, mechanisms controlling maintenance of identity, and accompanying regulation of plasticity, are largely unexplored and unknown.

Here, we identify two novel functions of the co-repressor *Tle4* in both acquisition and maintenance of neuron subtype identity of corticothalamic projection neurons. Embryonically, *Tle4* promotes acquisition of corticothalamic molecular and cellular identity, and blocks emergence of core characteristics of subcerebral / corticospinal projection neuron identity, including morphology, gene expression, axonal connectivity, and circuitry. Postnatally, *Tle4* is required to maintain corticothalamic molecular and projection identity during circuit maturation, avoiding potentially disruptive plasticity, but also limiting potentially beneficial plasticity.

We identify an epigenetic mechanism by which TLE4 controls the activation state of loci regulating the level of *Fezf2* expression by corticothalamic neurons during embryonic and postnatal development. This mechanism contributes importantly to distinction of cortical output (corticofugal) subtypes, and ensures appropriate maturation and maintenance of CThPN.

**Highlights:**

- *Tle4* promotes CThPN identity and blocks SCPN identity in early-born cortical neurons
- *Tle4* is necessary to maintain CThPN identity during circuit maturation
- TLE4-FEZF2 complex epigenetically regulates *Fezf2* expression in developing CThPN
- TLE4-FEZF2 regulates corticofugal subtypes distinction and maturation of CThPN

## Introduction

During embryonic development, genetic programs regulate acquisition and progressive refinement of distinct characteristics that constitute neuron identities, including gene expression, afferent and efferent connectivity, electrophysiological properties, molecular signaling, etc.(Greig et al., 2013; Lodato and Arlotta, 2015; Galazo et al., 2016). In the cerebral cortex, neuron subtype identity has been identified to be controlled both at progenitor stage and post-mitotically soon after terminal division (Frantz and Mcconnell, 1996; Shen et al., 2006), though multiple studies have revealed that aspects of identity remain malleable for at least the early period of postmitotic development (Rouaux et al., 2012). After initial differentiation, neuron identity appears actively maintained to preserve neuron subtype characteristics (Deneris and Hobert, 2014). In cortical neurons, some aspects of identity can be altered postnatally by genetic manipulation (Rouaux and Arlotta, 2013; De La Rossa et al., 2013), or changes in cortical input activity (Chevee et al., 2018). Some core features of neuron subtype identity, including gene expression, axonal projection, and electrophysiological properties, have been reprogrammed in embryonic and postnatal cortical neurons, resulting in the adoption of features of other neuron subtypes (Rouaux and Arlotta, 2013; De La Rossa et al., 2013). This raises important questions about how neuron subtypes maintain their identities in the dynamic context of the developing and mature cortex, and whether, e.g., identity might be regulated toward beneficial plasticity, regeneration, and restoration of functional circuitry.

Maintenance of neuronal subtype identity is important well beyond early development and organization of the nervous system. Failure of neuron identity maintenance has been implicated in emergence of neurodegenerative and psychiatric disorders later in life (Deneris and Hobert, 2014; Fazel Darbandi et al., 2018). Investigating how distinct cortical neurons acquire and maintain their identities is critical both to understand cortical development and dysgenesis, and to reveal intrinsic vulnerabilities of specific neurons to disorders. Further, better understanding of mechanisms controlling identity acquisition and maintenance *in vivo* could be instrumental to improve *in vitro* differentiation of clinically-relevant neuron subtypes for disease modeling. Beyond all of these areas, manipulation of identity maintenance might provide useful regulation of cellular plasticity toward neuronal subtype regeneration, repopulation, and/or other functional reconstitution in settings of neuronal circuit injury or compromise.

Here, we investigate neuron subtype-specific identity acquisition and maintenance by corticothalamic projection neurons (CThPN) during murine cortical development and maturation, in particular the function of *Tle4* in these processes. CThPN belong to the broader class of corticofugal projection neurons (neocortical output neurons), which include CThPN and subcerebral projection neurons (SCPN). CThPN are born early during corticogenesis, at embryonic days (E) 11.5-12.5 in mice, most of them reside in cortical layer VI, and they project to the thalamus. SCPN are born during mid-corticogenesis (peak at E13.5), reside in layer V, and project to the brainstem and spinal cord.

*Tle4* is a non-DNA binding transcriptional co-repressor with important functions in cell fate specification and differentiation in multiple cell types (Xing et al., 2018; Wheat et al., 2014; Yao et al., 1998). In the developing cortex, *Tle4* is frequently used as a corticothalamic “marker” gene (Molyneaux et al., 2015), and recently, it has been identified as an important regulator of the development of CThPN identity (Tsyporin et al., 2021). Previously, we identified *Tle4* as a “CThPN identity gene” since it is highly and specifically expressed by CThPN, not only during embryonic development, but also throughout early postnatal differentiation (Galazo et al., 2016). Because of its specific and temporally extended expression by CThPN, we hypothesized that *Tle4* might function combinatorially with other central regulators of corticofugal and CThPN subtype development to regulate acquisition and stability of CThPN identity during development, circuit formation, and maturation.

Our work presented here identifies that *Tle4* is necessary for both acquisition and maintenance of CThPN identity. In the absence of *Tle4*, CThPN do not develop normal molecular, morphological, or projection identity, and instead acquire hallmark characteristics of SCPN. Importantly, *Tle4* is also necessary during postnatal maturation to maintain stable CThPN identity. Loss of *Tle4* function at differentiated, relatively mature stages, when CThPN have already established thalamic connections and circuitry, results in upregulation of genes characteristic of SCPN, with extension of aberrant axonal projections to subcerebral targets. We identify an epigenetic mechanism mediated by TLE4 that regulates *Fezf2* expression level during acquisition and maintenance of CThPN identity. This mechanism functions to block the emergence of SCPN identity during CThPN specification and maturation, then maintain the distinct identity of CThPN. This work thus reveals a critical function of *Tle4* not only regarding CThPN, but more broadly in the differentiation and continued distinction of the two main classes of corticofugal neurons– CThPN and SCPN. *Tle4* first controls CThPN vs. SCPN developmental distinction, then maintains stable and sharply bounded CThPN identity during maturation. In this way, *Tle4* actively limits plasticity between these developmentally related classes of cortical projection neurons.

## Results

### *Tle4* is expressed by developing and mature CThPN

Previous studies have shown *Tle4* is expressed in the embryonic cortex, and postnatally in layer VI (Yao et al., 1998; Allen and Lobe, 1999), where it is specifically expressed by CThPN (Galazo et al., 2016; Molyneaux et al., 2015; Tsyporin et al., 2021). *Tle4* specificity to layer VI-CThPN has been determined during early postnatal development (Tsyporin et al., 2021), however, whether *Tle4* neuron-subtype specific expression varies over time, and to what extent *Tle4* is expressed by distinct subsets of CThPN has not been investigated.

Using *in situ* hybridization (ISH), we find that *Tle4* is first expressed in the cortical plate at E12.5 by the earliest born cortical neurons (developing CThPN). Expression by progenitors is not detected. Embryonically, *Tle4* is strongly expressed in the subplate and deep cortical plate, where developing CThPN are located (Fig. 1). In previous work comparing the gene expression profiles of distinct cortical projection neurons at embryonic and early postnatal stages (Galazo et al., 2016), we already determined that *Tle4* expressing cells in the cortical plate at E18, and layer VI at postnatal day (P) 3 and P6 are CThPN.

**Figure 1.**
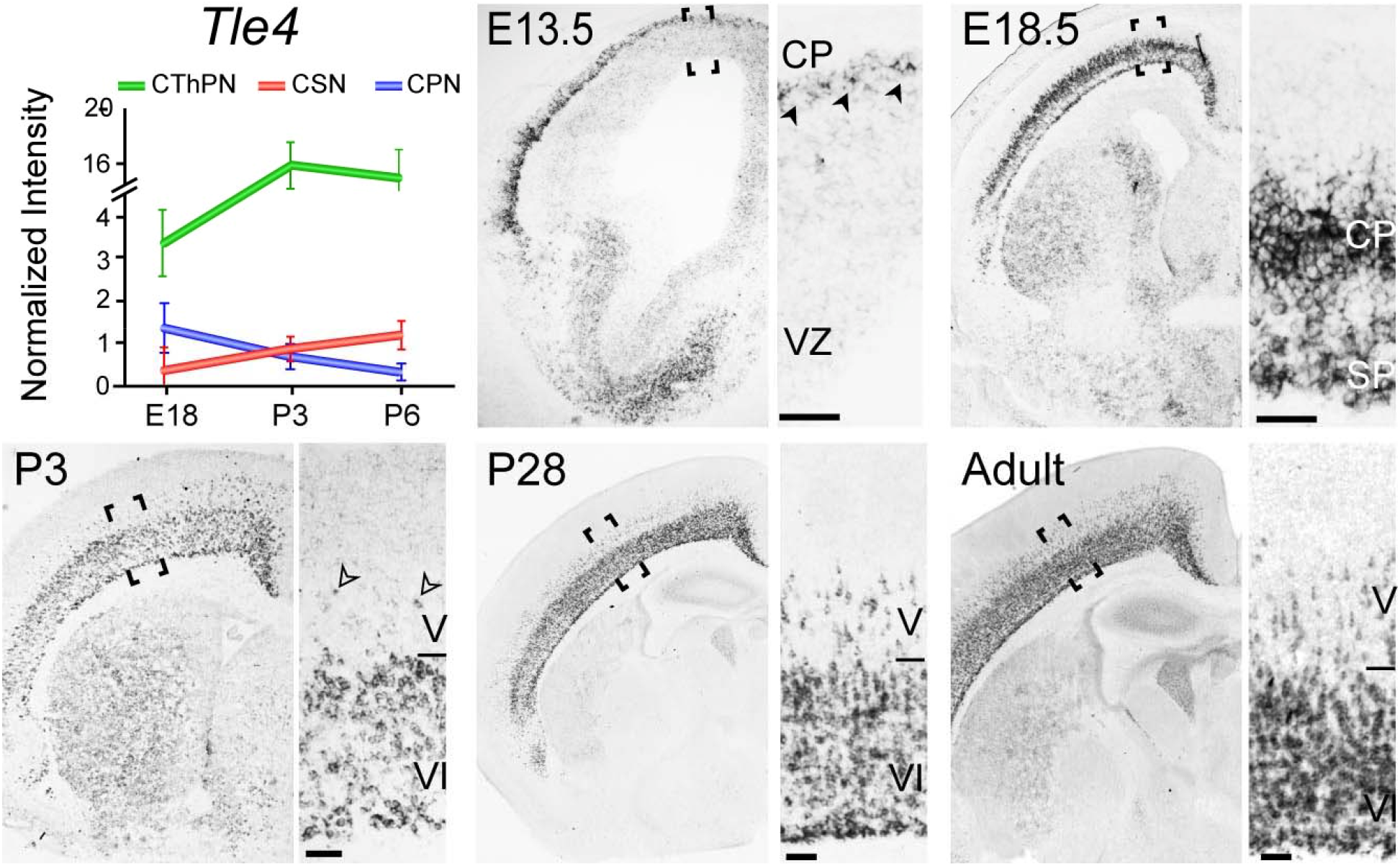
Developmental dynamics of *Tle4* expression. Temporal profile of *Tle4* expression by CThPN (green), CSN (red), and CPN (blue) from microarray analysis at E18, P3, and P6 (adapted from Galazo et al., 2016). *In situ* hybridization reveals that *Tle4* is first expressed by newly-postmitotic neurons in the cortical plate by E13.5 (black arrowheads). Expression increases through embryonic development, shown at E18.5. Postnatally, strong expression continues in layer VI from the neonatal period, shown at P3, through maturity, shown at P28. At P3, almost all *Tle4^+^* neurons are in layer VI, with a few in layer V (open arrowheads). At P28, the predominant population is still in layer VI, but modestly more neurons in layer V express *Tle4*. Scale bars, 50 μm.

During the first postnatal week, most *Tle4*-expressing neurons reside in layer VI, while a smaller proportion resides in layer V. *Tle4* expression in layer V increases during postnatal development and persists in adults (Fig. 1; compare P3, P28, Adult). *Tle4* remains highly expressed in layer VI throughout postnatal maturation and in adults.To determine the neuron subtype identity of TLE4^+^ neurons in both, layer VI and layer V, we retrogradely labeled three major projection neuron subtypes– CThPN, SCPN, and callosal projection neurons (CPN), and immunolabeled for TLE4. We confirmed that essentially all TLE4^+^ cells in layer VI are CThPN by retrograde labeling from the thalamus with Cholera toxin-b (Ctb^+^) or Fast blue (Fb^+^) (Supplementary Fig. 1a), and by co-labeling with a layer VI-CThPN reporter (*Ntsr1-Cre;tdTomato^fl^*) (Tasic et al., 2016) (Supplementary Fig. 1d). A small fraction of the CThPN population resides in layer V. Layer V-CThPN are functionally different from layer VI-CThPN, and send dual projections to thalamus and brainstem (Guillery and Sherman, 2002). To investigate whether TLE4^+^ neurons in layer V are Layer V-CThPN, we performed dual retrograde labeling from thalamus (Fb) and cerebral peduncle (Ctb). We find that 62% of the TLE4^+^ neurons in layer V have dual projections to thalamus and brainstem (double-labeled Fb^+^-Ctb^+^) (Supplementary Fig. 1b). This percentage likely underestimates the number of TLE4^+^ neurons identified as layer V-CThPN, since dual retrograde labeling usually captures a subtotal fraction of the population. Strikingly, only 2.5% of the TLE4^+^ neurons in layer V are SCPN (documented projection to brainstem) without a projection to the thalamus (Supplementary Fig. 1b). We also studied whether some TLE4^+^ neurons in layer V are CPN. We find that only 1.8% of labeled CPN are TLE4^+^ (1.4% are layer V-CPN, and 0.4% are layer VI-CPN, Supplementary Fig. 1c). These experiments reveal that essentially all layer V-TLE4^+^ neurons are CThPN.

Together, these data indicate that *Tle4* is constitutively and specifically expressed by CThPN, regardless of their layer location or CThPN-functional subtype.

### *Tle4* promotes differentiation of CThPN identity, and prevents acquisition of SCPN identity in early-born cortical neurons

*Tle4* is a corepressor with important functions in cell fate determination and development of multiple tissues (Xing et al., 2018; Wheat et al., 2014; Yao et al., 1998). Function of *Tle4* in cortical development has started to be investigated; recently, one function of *Tle4* in CThPN differentiation has been revealed using a mutant allele with a *LacZ-ires-Plap* cassette in the 4^th^ intron of *Tle4* to disrupt its normal expression (Tsyporin et al., 2021). To potentially more broadly investigate functions of *Tle4* in cortical development, we employed a constitutive *Tle4* null mutant mouse allele referred to here as *Tle4^KO^*. This null allele disrupts transcription after exon 1 of *Tle4* and produces no functional TLE4 protein (Wheat et al., 2014).

Using *Tle4^KO^* mice for loss-of-function, we investigated function(s) of *Tle4* in CThPN differentiation. As a first step, we analyzed cortical cytoarchitecture in *Tle4^KO^* mice at P8. There is no difference in cortical thickness or in cytoarchitecture of the superficial-layers between wildtype control and *Tle4^KO^* cortices (Fig. 2a). Strikingly, however, *Tle4^KO^* layer V is significantly expanded at the expense of layer VI. Further, there is a striking increase in the number of large pyramidal neurons (>20 μm diameter), normally only abundant in layer V, at cortical depths that normally correspond to layer VI (neurons > 20 μm diameter in layer VI: *Tle4^KO^* 42% vs. 18% wildtype control, p<0.01; Fig. 2a). This aberrant presence in layer VI of these largest diameter neurons is not due to incomplete migration of layer V neurons at P8, since it persists in adult *Tle4^KO^* mice (Supplementary Fig. 2a). This striking layer VI reduction with layer V expansion in *Tle4^KO^* mice hypothetically might be due to abnormal production, migration, and/or differentiation of neurons. To distinguish between these possibilities, we performed BrdU birth-dating at E11.5, E12.5, E13.5, and E14.5 (peaks of subplate, CThPN, SCPN, and superficial layer-CPN generation, respectively), and analyzed the number and distribution of BrdU^+^ neurons across layers at P6. There is no significant difference in the number or distribution of BrdU^+^ neurons labeled at any of these stages (Supplementary Fig 2b), indicating that neuronal generation and migration are unaffected in *Tle4^KO^* mice. These results are consistent with the recent report of normal neuron generation and migration in the intron 4 insertion *Tle4^Laz^* mutant mice (Tsyporin et al., 2021), and indicate *Tle4* regulation of layer VI-V neuronal differentiation.

**Figure 2.**
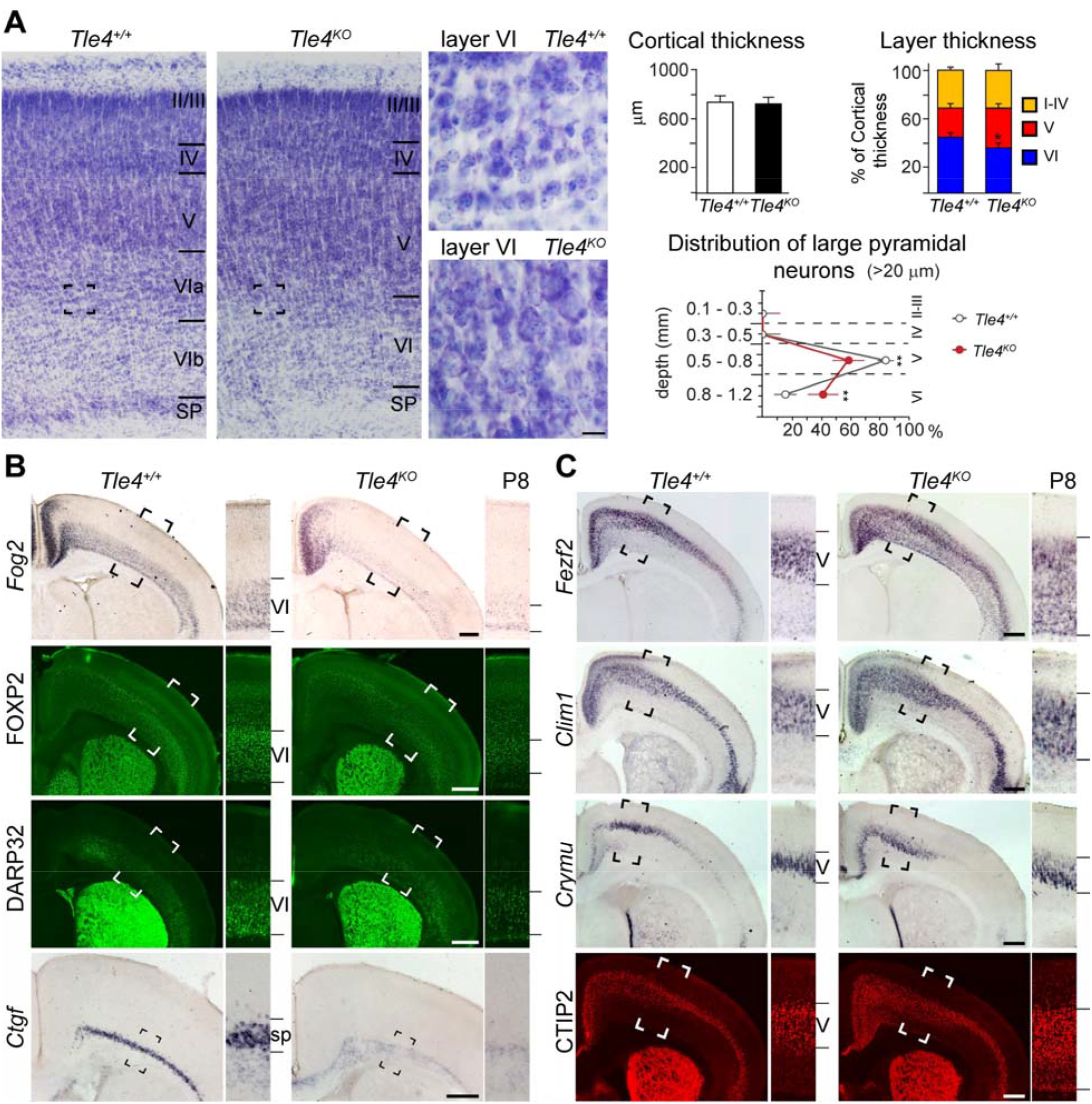
In the absence of *Tle4* function, CThPN do not develop normal identity, and acquire cardinal somatic morphology and gene expression characteristic of SCPN. (A) Nissl straining of cortex in wildtype and *Tle4^KO^* reveals cytoarchitectural abnormalities in *Tle4^KO^* cortex. Insets show at high magnification the somatic morphology of neurons at depth corresponding to layer VIa in wildtype and *Tle4^KO^* cortex. There is no difference in overall cortical thickness between *Tle4^KO^* and wildtype controls, but layer V is substantially expanded, and layer VI is substantially reduced, in *Tle4^KO^* cortex. Distribution of large pyramidal neurons with diameter >20 μm at a range of cortical depths from layer I. In wildtype cortex, 82% of large pyramidal neurons reside in layer V and only 18% in layer VI, while in *Tle4^KO^* mice, only 58% of large pyramidal neurons reside in layer V, with a substantially increased 42% in layer VI. (B-C) In the absence of *Tle4*, expression of CThPN control genes and markers *Fog2*, FOXP2, TBR1, and *Ctgf* decrease, and expression of SCPN control genes and markers *Fezf2, Clim1, Crymu*, and CTIP2 increase at P8. Insets magnify the boxed areas. Scale bars, (a) 20 μm, (b-c) 500 μm.

Our data showing the presence of large diameter pyramidal neurons– typically limited to layer V– in what would normally be layer VI indicates that early-born neurons that would normally become smaller pyramidal CThPN do not differentiate normally in *Tle4^KO^* mice; they acquire at least cardinal size and morphological features of SCPN. Recent work by Tsyporin et al., 2021 using the *Tle4^Laz^* mutant allele reveals important functions of *Tle4* in regulating basal dendritic branching, excitability, and molecular identity of CThPN (Tsyporin et al., 2021). Beyond both of these sets of results, we investigated whether other aspects of CThPN identity are altered, and in particular whether additional morphological and molecullar characteristics of SCPN identity emerge in early-born neurons positioned in layer VI in *Tle4^KO^* cortex. Long apical dendrites with tufts reaching layers II-III and layer I are characteristic of SCPN, while CThPN usually have shorter apical dendrites extending only to layer IV. We visualized dendrites of what are normally CThPN using the normally CThPN-specific reporter *Ntsr1-Cre:tdTomato^fl^*. We find many dendritic apical tufts aberrantly extending into superficial layers II/III and I in *Tle4^KO^* cortex, but not in wildtype controls (Supplementary Fig. 2c). Next, we investigated whether CThPN molecular identity is altered in *Tle4^KO^* mice, using *in situ* hybridization and immunolabeling for known CThPN molecular controls and markers (Galazo et al., 2016). There is very substantial downregulation of *Ctgf,* DARP32, *Fog2,* FOXP2, NURR1, and *Shb* by these abnormal *Tle4^KO^* layer VI neurons compared to wildtype, and moderate downregulation of TBR1 and *Cxxc5* (Fig. 2b, and Supplementary Fig. 2d). Importantly, and consistently with the expression data reported in *Tle4^Laz^* mutant mice, we find strong upregulation of SCPN genes (*Fezf2*, CTIP2, *Clim1, Crymu, Bhlhb5, Id2, Pcp4,* and *S100a10*) by the aberrant layer VI neurons in *Tle4^KO^* cortex (Fig. 2c, and Supplementary Fig. 2e). Of particular note, there is especially strong upregulation across layer VI of *Fezf2,* known as an SCPN selector gene (Fig. 2c). Together, these results indicate that early-born neurons do not develop appropriate CThPN molecular identity or morphology in *Tle4^KO^* cortex, and instead acquire central SCPN molecular identity and morphology– suggesting either an improper identity refinement between the subtypes or a fate switch between the subtypes. SCPN identity would be even further confirmed if the quintessential, defining SCPN characteristic– subcerebral axon projections– are formed by these fate-switched SCPN (which normally would have been CThPN).

To investigate whether abnormally developing CThPN in *Tle4^KO^* mutants switch their fate to become SCPN, we directly investigated whether they extend subcerebral axonal projections. We labeled SCPN by injecting Ctb into the cerebral peduncle at P3 and determined their location in the cortex at P6. In wildtype controls, Ctb^+^ SCPN are restricted to layer V. In *Tle4^KO^* mice, we find significanly more Ctb^+^ SCPN, and they are located in layer V, layer VI, and subplate. Many of the additional Ctb^+^ SCPN are located just deep to the normal layer V border, giving the appearance of a thicker layer V that extends down into layer VI. We refer to this downward extension of the layer V as ‘extended layer V’ (V_ex_) (Fig. 3a, c). Combining Brdu-birthdating and SCPN retrograde labeling, we find significantly more SCPN (Ctb^+^) born during the usual peak of CThPN neurogenesis in *Tle4^KO^* cortex (BrdU^+^-Ctb^+^/Ctb^+^ in *Tle4^KO^* 11% vs. 0% in widtype control, BrdU at E11.5, p<0.01; BrdU^+^-Ctb^+^/Ctb^+^ *Tle4^KO^* 8% vs. controls 1%, BrdU at E12.5, p<0.01, Fig. 3c). The percentage of neurons born at E12.5 that strongly express CTIP2 is consistently increased in *Tle4^KO^* cortex (BrdU^+^-CTIP2^+^/BrdU^+^ in *Tle4^KO^* 42% vs. wildtype control 21%, BrdU at E12.5, p<0.01, Fig. 3c), and the percentage of neurons born at E11.5-E12.5 that express FOG2 is decreased in *Tle4^KO^* cortex (BrdU^+^-FOG2^+^/BrdU^+^ *Tle4^KO^* 8% vs. control 37%, BrdU at E11.5, p<0.01; *Tle4^KO^* 10% vs. control 28%, BrdU at E12.5, p<0.01) (Fig. 3d). These results strongly indicate that, in *Tle4^KO^* mice, a substantial number of early-born neurons that normally become CThPN fate-switch to become SCPN.

**Figure 3.**
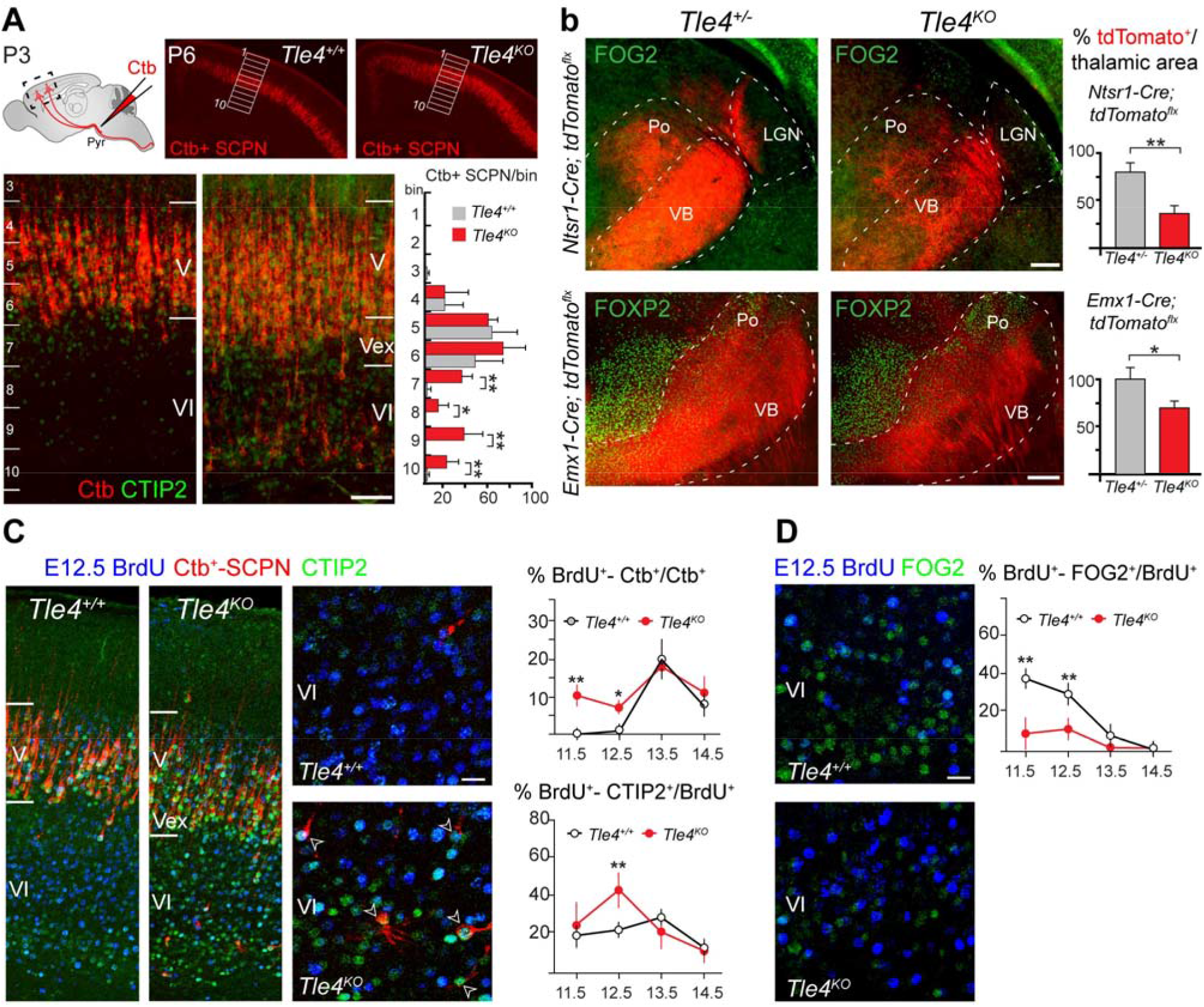
In *Tle4^KO^* cortex, more early born neurons become SCPN and fewer develop as CThPN. (**A**) Substantially more neurons are retrogradely labeled by injection of Ctb into the cerebral peduncle in *Tle4^KO^* than in wildtype controls. Schematic of Ctb injections performed at P3, and labeling assessed in the cortex at P6. (**B**) Axonal projections to the thalamus are substantially reduced in *Tle4^KO^* mice compared to controls. CThPN axons are labeled by tdTomato endogenous fluorescence from the CThPN-specific reporter *Ntsr1-Cre:tdTomato^fl^* and the cortex-specific reporter *Emx1-Cre:tdTomato^fl^*. Quantification of tdTomato^+^-pixel density in somatosensory thalamic nuclei, posterior (Po) and ventral posteromedial (VPm) nuclei, in *Ntsr1-Cre:tdTomato^fl^:Tle4^+/−^*, *Ntsr1-Cre:tdTomato^fl^:Tle4^KO^*, *Emx1-Cre:tdTomato^fl^:Tle4^+/−^*, and *Emx1-Cre:tdTomato^fl^:Tle4^KO^* mice performed at P6. Quantification performed in the somatosensory thalamic nuclei, posterior (Po) and ventrobasal (VB) nuclei. The lateral geniculate (LGN) nucleus labeled for reference. Values expressed as percentage normalized to controls. Two-tailed t-test was used (* p<0.05; ** p<0.01). (**C-D**) More early born neurons in *Tle4^KO^* cortex project as SCPN and express CTIP2 strongly, and fewer express FOG2, compared to wildtypes. (**c**) Quantification of SCPN, identified by retrograde labeling from the cerebral peduncle, labeled by BrdU at E11.5, E12.5, E13.5, and E14.5. Significantly more neurons born at E11.5, E12.5 are SCPN in *Tle4^KO^* mice compared to wildtypes. Significantly more neurons born at E12.5 strongly express CTIP2 in *Tle4^KO^* compared to wildtypes. Retrograde labeling performed at P3, and analysis performed at P6. (**D**) Significantly fewer neurons born at E11.5 and E12.5 express the CThPN molecular control FOG2 in *Tle4^KO^* mice compared to wildtypes. ANOVA test (* p<0.05; ** p<0.01). Scale bars, (a) 100 μm, (b) 200 μm, (c-d) 20 μm.

Early born neurons might still develop thalamic projections despite their abnormal CThPN molecular identity. To investigate this hypothesis, we visualized CThPN axons in the thalamus using *Ntsr1-Cre:tdTomato^fl^* reporter (CThPN-specific) and *Emx1-Cre:tdTomato^fl^* (cortex-specific) reporter, crossed into the *Tle4^KO^* background. We find a strong reduction of thalamic projections in *Ntsr1-Cre:tdTomato^fl^:Tle4^KO^* mice (0.45-fold reduction, p<0.01, Fig. 3b), and in *Emx1-Cre:tdTomato^fl^:Tle4^KO^* mice (0.7-fold reduction, p<0.05, Fig. 3b) compared to their respective *Ntsr1-Cre:tdTomato^fl^:Tle4^+/−^* and *Emx1-Cre:tdTomato^fl^:Tle4^+/−^* controls. Although thalamic projections in *Ntsr1-Cre:tdTomato^fl^*:*Tle4^KO^* and *Emx1-Cre:tdTomato^fl^:Tle4^KO^* are reduced, the innervation pattern seems grossly normal. Therefore, CThPN still innervate the thalamus in the mutants. We analyzed whether layer VI fate-converted SCPN also bear a projection to thalamus, which would reflect their original corticothalamic fate. We performed dual retrograde labeling from thalamus and brainstem in mutants and controls. In both *Tle4^KO^* and wildtype controls, we only find neurons with dual projections in layer V (normal layer V-CThPN population) (Supplementary Fig. 3), suggesting that fate-converted SCPN override their initial corticothalamic projection identity. Together, these results demonstrate that in *Tle4^KO^* cortex, early-born neurons do not acquire normal CThPN identity, and while many still project to the thalamus, others fate-switch and acquire the morphological, molecular, and projection identity typical of SCPN.

### *Fezf2, Ctip2, and Fog2* are dysregulated in all fate-converted SCPN in *Tle4^KO^* mice

In *Tle4^KO^* mice, all early-born neurons develop abnormal molecular identity, however, only a subset becomes SCPN (fate-converted SCPN). To investigate the molecular changes specifically associated with this switch of projection identity, we identified fate-converted SCPN by retrograde labeling from the brainstem and analyzed expression of genes critical for axonal projection identity in SCPN (*Fezf2*, *Ctip2*, and *Bhlbh5*) (Molyneaux et al., 2005; Chen et al., 2005a; Chen et al., 2005b; Arlotta et al., 2005; Joshi et al., 2008), and CThPN (*Tbr1, Fog2*) (Hevner et al., 2001; Mckenna et al., 2011; Han et al., 2011; Galazo et al., 2016).

*Fezf2* is critical for SCPN development (Molyneaux et al., 2005; Chen et al., 2005a; Chen et al., 2005b). In *Tle4^KO^* mice, *Fezf2* is strongly upregulated by virtually all fate-converted SCPN (99% of Ctb^+^ in layer ‘V_ex_’, 95% of Ctb^+^ layer VI and subplate), and also by layer VI neurons that do not become SCPN (Fig. 4a, f). This suggests that, while *Fezf2* upregulation is required to convert early-born neurons into SCPN, is not the only factor responsible for the change of the projection pattern. CTIP2 upregulation is less widespread than *Fezf2* upregulation, however, virtually all fate-converted SCPN strongly express CTIP2 (97% of layer ‘V_ex_’ SCPN, 96% of layer VI-subplate SCPN, Fig. 4b, f). In contrast, *Bhlhb5* is specifically expressed by fate-converted SCPN in layer ‘V_ex_’, but not by fate-converted SCPN in layer VI (93% in ‘V_ex_’ SCPN, <1% in layer VI-subplate SCPN, Fig. 4c, f). *Bhlhb5* is critical for the differentiation of two specific SCPN subsets, the caudal corticospinal motor neurons, and the corticotectal projection neurons (Joshi et al., 2008). Differential expression of SCPN genes in distinct fate-converted SCPN subsets suggests differences in their projection pattern.

**Figure 4.**
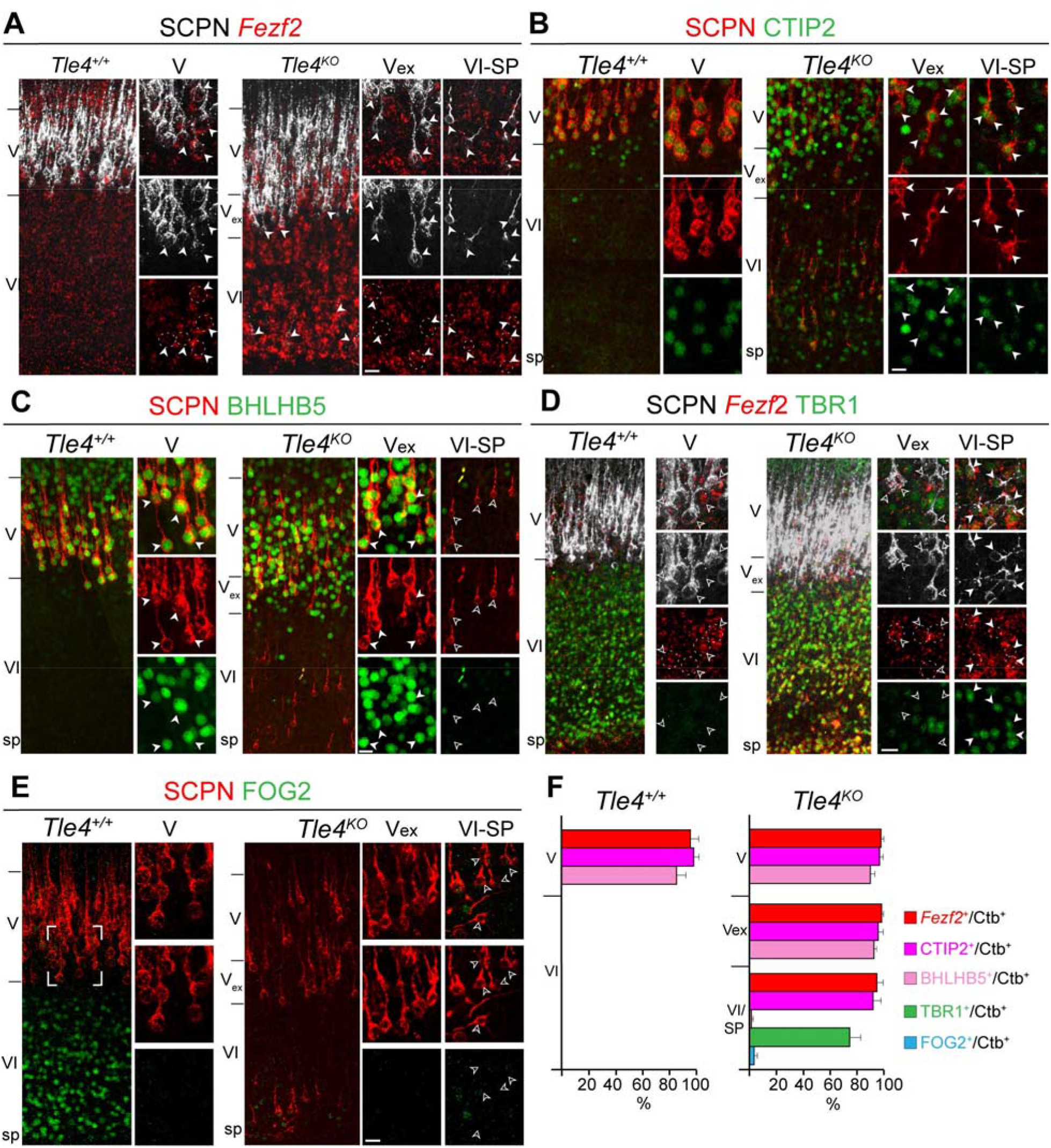
Expression of central SCPN and CThPN molecular controls by fate-converted SCPN in *Tle4^KO^* mice. (**A**) Most neurons in the deep layers express *Fezf2* strongly in *Tle4^KO^* cortex, with aberrantly strong layer VI expression. ISH for *Fezf2* (red), and SCPN retrograde labeling with Ctb (white), reveals that virtually all SCPN express *Fezf2* (white arrowheads), regardless of their location in layer V, layer V_ex_, layer VI, or subplate. *Fezf2* is also expressed at high level in layer VI of *Tle4^KO^* mice by neurons that do not become SCPN. (**B**) Immunolabeling for CTIP2 (green), and SCPN retrograde labeling with Ctb (red), reveals that virtually all SCPN express CTIP2 in *Tle4^KO^* mice, regardless of their location in layer V, layer V_ex_, layer VI, or subplate (white arrowheads). (**C**) Immunolabeling for BHLHB5 (green), and SCPN retrograde labeling with Ctb (red), reveals that BHLHB5 is expressed in *Tle4^KO^* mice by fate-converted SCPN in layer V_ex_ (white arrowheads), but not by fate-converted SCPN in layer VI or subplate (open arrowheads). (**D**) Immunolabeling for TBR1 (green), ISH for *Fezf2* (red), and SCPN retrograde labeling with Ctb (white) reveal that fate-converted SCPN in layer V_ex_ express *Fezf2* but not TBR1 (open arrowheads). In contrast, fate-converted SCPN in layer VI express both *Fezf2* and TBR1 (solid arrowheads). Dashed lines encircle *Fezf2^+^* neuron somata to facilitate visualization. (**E**) Immunolabeling for FOG2 (green), and SCPN retrograde labeling with Ctb (red), reveals that barely any fate-converted SCPN in layer VI or subplate express FOG2 (open arrowheads). (**F**) Percentage of SCPN expressing *Fezf2*, CTIP2, BHLHB5, TBR1 or FOG2 in layer V and layer VI-subplate in wildtype mice, and in layer V, layer V_ex_, and layer VI-subplate in *Tle4^KO^* mice. Scale bars, 20 μm (a-e).

In *Tle4^KO^* mice, TBR1, which normally strongly determines CThPN identity and represses *Fezf2* in early-born neurons (Hevner et al., 2001; Han et al., 2011), is slightly reduced, but expression remains in layer VI and subplate neurons, where it is co-expressed with *Fezf2*. TBR1 is not expressed by fate-converted SCPN in layer ‘V_ex_’, but it is expressed by fate-converted SCPN in layer VI (Fig. 4d, f). This result indicates that the level of TBR1 repression of *Fezf2* is context-dependent, and that TBR1 expression does not necessarily preclude axonal projection to subcerebral targets.

Importantly, FOG2 is strongly downregulated in *Tle4^KO^* mice, and is barely expressed by any fate-converted SCPN (0% FOG^+^-Ctb^+^ in ‘V_ex_’, 3% FOG^+^-Ctb^+^ in layer VI-subplate, Fig. 4e, f). FOG2 downregulates *Ctip2* expression in a context-specific manner and reduces axonal projections to the brainstem when mis-expressed in SCPN (Galazo et al., 2016). Collectively, these data reveal that expression of *Fezf2, Ctip2,* and *Fog2* are dysregulated by all fate-converted SCPN in *Tle4^KO^* mice, and that fate-converted SCPN in layer ‘V_ex_’ are more similar to normal wildtype SCPN population than are the fate-converted SCPN in layer VI.

### Area-specific projection identity is preserved in the absence of *Tle4*

In the context of the combined morphological, molecular, and projection shifts presented above, it remained possible that, while fate-converted SCPN project into the cerebral peduncle, they might not reach the more precise, area-appropriate targets in the brainstem and spinal cord corresponding to their cortical area of origin, and project instead to inappropriate targets. To investigate this possibility, first, we investigated whether there are ectopic subcerebral projections in *Tle4^KO^* brains. In *Tle4^KO^* mice, we identified fate-converted SCPN across cortical areas via retrograde labeling from the cerebral peduncle. Since all SCPN, including fate-converted SCPN, strongly express *Fezf2*, we used *Fezf2*^+*/PLAP*^ mice as a reporter (Chen et al., 2005a) to label SCPN axonal projections in the brainstem. In P21 *Fezf2*^+*/PLAP*^*:Tle4*^+/−^ mice, PLAP-positive axons appropriately innervate the superior colliculus and extend along the pyramidal tract. In *Fezf2*^+*/PLAP*^:*Tle4*^KO^ mice, there are more PLAP-positive axons in the superior colliculus, cerebral peduncle, and pyramidal tract, but no ectopic projections to abnormal targets (Supplementary Fig. 4a,b, and Supplementary Fig. 5a). The same is true using *Emx1-Cre:tdTomato^fl^* reporter mice to label all axons from cortical projection neurons, regardless of their expression of *Fezf2*. There are more tdTomato^+^ axons in the cerebral peduncle and in the superior colliculus in *Emx1-Cre:tdTomato^fl^:Tle4*^KO^ compared to control mice, but no frankly ectopic projections (Supplementary Fig. 5b), indicating that fate-converted SCPN do not project aberrantly to inappropriate targets.

Next, we investigated whether fate-converted SCPN project with the same area-specificity as normal SCPN. In adult wildtype mice, SCPN in somatosensory cortex project to the pons and superior colliculus, while SCPN in visual cortex project only to the superior colliculus (Greig et al., 2013; Lodato and Arlotta, 2015). Using double labeling from the pyramidal tract at the level of the pons (Ctb) and superior colliculus (Fb), we investigated whether fate-converted SCPN preserve appropriate area-specific projection targeting. As expected in adult wildtype controls, there are SCPN in layer V of somatosensory cortex double-labeled from pons and superior colliculus (Ctb^+^-Fb^+^) (Fig. 5a), but SCPN in layer V of visual cortex are labeled from only superior colliculus (Fb^+^) (Fig. 5b). In *Tle4^KO^* mice, similarly, there are Ctb^+^-Fb^+^ SCPN in somatosensory cortex, but they are present more broadly in layers V, V_ex_, and VI (Fig. 5a). Importantly, in *Tle4^KO^* visual cortex, there are only Fb^+^ SCPN, but again located more broadly in layer V, V_ex_, and VI (Fig. 5b), indicating that fate-converted SCPN remain area-appropriate. In particular, in visual cortex, SCPN project specifically to their corresponding visual targets, but not to the pons, as do SCPN from other areas. We investigated area-specificity further, and confirmed area-specificity of *Tle4^KO^* fate-converted SCPN projections to the spinal cord. While there are more corticospinal neurons innervating the cervical spinal cord (CSNc) in *Tle4^KO^* mice compared to wildtype controls, in both groups, CSNc are restricted to rostral motor cortex (Fig. 5c). Together, these results indicate that, while *Tle4* is critical for the appropriate differentiation and distinction of CThPN and SCPN subtype identities, it is not necessary for acquisition of area-specific identity. Fate-converted SCPN resulting from loss of *Tle4* function acquire area-appropriate and area-specific axonal connectivity matching that of endogenous SCPN.

**Figure 5.**
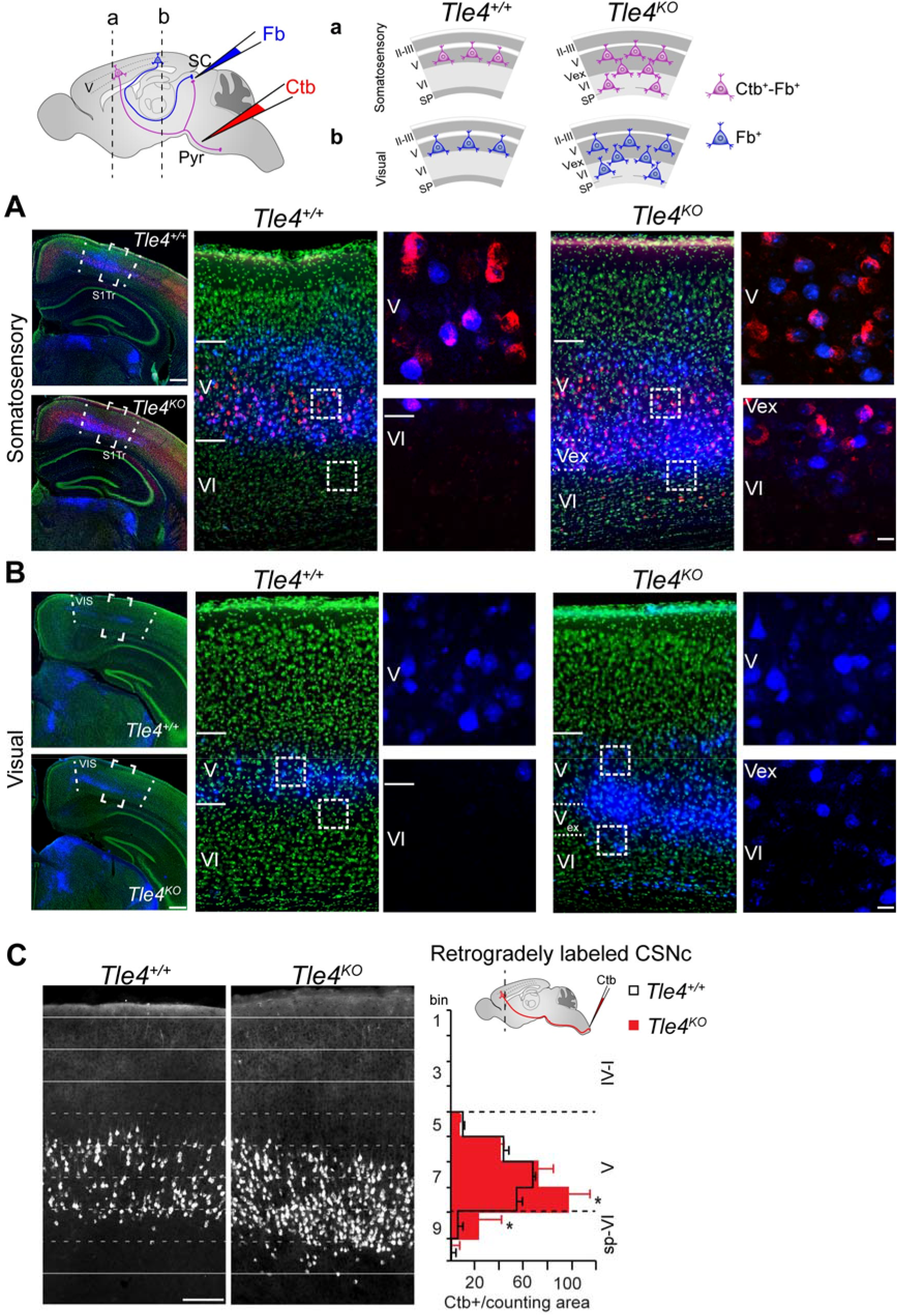
Area-specific projection identity is preserved in the absence of *Tle4*. (**A-B**) Dual retrograde labeling from the superior colliculus and pons to reveal area-specific projection patterns in the Somatosensory and Visual cortices to these targets. Fb was injected into the superior colliculus (SC) and Ctb was injected into the pyramidal tract (Pyr) in adult *Tle4^KO^* and control mice. Top, schematics of the dual retrograde approach, and retrograde labeling patterns observed in control and *Tle4^KO^* cortices. (**A**) In controls, double-retrogradely labeled neurons (Ctb^+^-Fb^+^) are located in the layer V the somatosensory cortex, but not in the visual cortex. In *Tle4^KO^,* Ctb^+^-Fb^+^ are also found in the somatosensory cortex, but in layers V, ‘Vex’, and VI (insets). (**B**) No Ctb^+^-Fb^+^ neurons were found in the visual cortex, either in control or *Tle4^KO^* mice. Fb^+^ are located in layer V of wildtype mice, and in layer V, ‘Vex’, and VI in *Tle4^KO^* mice. High magnification images correspond to squared areas. Primary somatosensory trunk area (S1Tr), Visual cortex (VIS). (**C**) More corticospinal motor neurons are labeled in rostral motor cortex by injection of Ctb into the corticospinal tract at the level of cervical spinal cord in *Tle4^KO^* mice compared to controls. Retrograde labeled corticospinal neurons innervating the cervical spinal cord (CSNc) are restricted to the rostral motor cortex in *Tle4^KO^* and controls. Right panel shows schematic of Ctb injections performed at P3 and quantification of Ctb^+^ CSNc performed in the rostral motor cortex at P14. For quantification of Ctb^+^ CSNc, the rostral motor cortex was divided in 10 equal bins, spanning the depth of the cortex. Bins 1-4 correspond to layers I-IV, bins 5-8 correspond to layer V, and bins 9-10 correspond to layer VI and subplate. Two-tailed t-test was used (* p<0.05). Scale bars, 500 μm (a-b, low magnification images), 20 μm (a-b, insets), 200 μm (c).

### *Tle4* is necessary to maintain CThPN identity during circuit maturation

Since *Tle4* is constitutively expressed by CThPN from development through adulthood (Fig. 1), we investigated whether *Tle4* might also function during postnatal CThPN maturation. We employed temporal conditional gene deletion using *Tle4^floxed^* mice (*Tle4^fl/fl^*) (Wheat et al., 2014). We induced recombination of the *Tle4^fl/fl^* allele at stages when CThPN axons are innervating thalamic nuclei (starting perinatally) (Jacobs et al., 2007; Grant et al., 2012), and when CThPN are forming synaptic connections with thalamic neurons (starting during the first postnatal week) (Golshani et al., 1998; Liu et al., 2004).

We investigated function of *Tle4* when CThPN axons are innervating the thalamus, employing conditional gene deletion in *Ntsr1-Cre:Tle4^fl/fl^* mice. First, we determined when *Cre* expression starts in *Ntsr1-Cre* mice, and its specificity to CThPN at perinatal stages. Robust *Cre* expression starts at E18-P0 in *Ntsr1-Cre* mice (Supplementary Fig. 6a), thus recombination of the *Tle4^fl/fl^* allele in *Ntsr1-Cre:Tle4^fl/fl^* mice will occur in that time frame. We also confirmed that *Cre* is expressed specifically by CThPN perinatally, as it is postnatally (Supplementary Fig. 6b).

In *Ntsr1-Cre:Tle4^fl/fl^* mice, loss of *Tle4* function causes upregulation of *Fezf2* and CTIP2 by CThPN, as in *Tle4^KO^* mice (Fig. 6a, b). To investigate whether CThPN convert into SCPN in *Ntsr1-Cre:Tle4^fl/fl^* mice, we retrogradely labeled SCPN at P3, and determined their location in the cortex at P7. In *Ntsr1-Cre:Tle4^fl/fl^* mice, in addition to the normal layer V SCPN, there are retrogradely labeled SCPN in layers V_ex_ and VI, though these are less abundant than in *Tle4^KO^* mice. In *Tle4^fl/fl^* controls, SCPN are restricted appropriately to layer V (Fig. 6c, d). In *Ntsr1-Cre:Tle4^fl/fl^* mice, layer V_ex_ SCPN and layer VI SCPN upregulate CTIP2 and downregulate FOG2 expression, as in *Tle4^KO^* mice (Fig. 6d).

**Figure 6.**
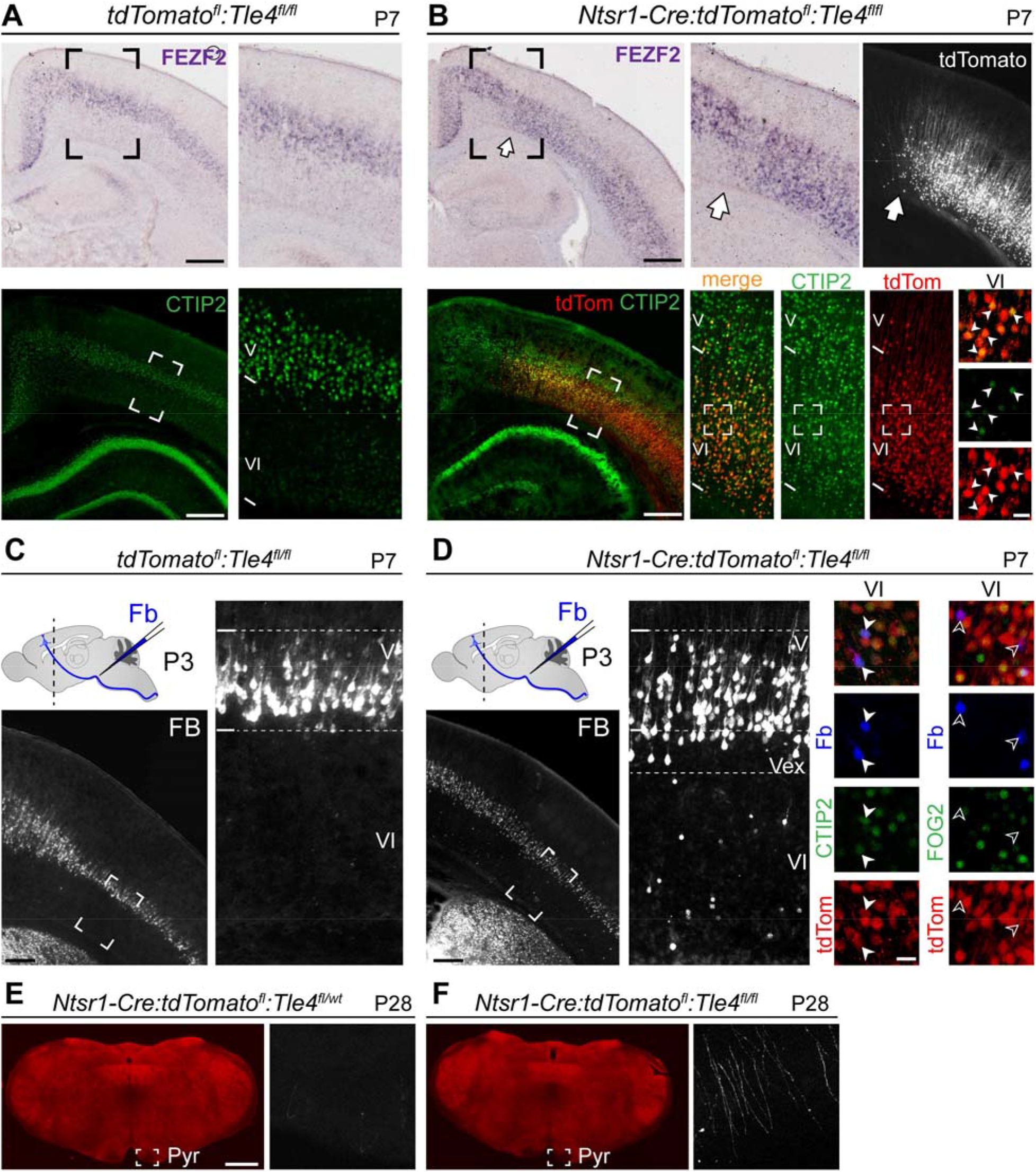
*Tle4* is required perinatally to maintain CThPN identity during thalamic innervation. (**A-B**) Perinatal loss of *Tle4* function results in upregulation of cardinal SCPN control genes. *Fezf2* ISH and CTIP2 immunolabeling reveal dramatic upregulation and spatial expansion in layer V_ex_ and VI of these SCPN molecular controls by Ntsr1-Cre-expressing neurons in *Ntsr1-Cre:tdTomato^fl^:Tle4^fl/fl^* mice. Higher magnification panels correspond to the boxed areas in the low magnification images. White arrow indicates the limit of Ntsr1-Cre expression, between somatosensory and motor cortices. (**C-D**) Retrograde labeling by injection of Fb into the cerebral peduncle at P3 reveals the presence of SCPN in layer VI and layer V_ex_ after loss of *Tle4* function in *Ntsr1-Cre:tdTomato^fl^:Tle4^fl/fl^* mice, but not in *tdTomato^fl^:Tle4^fl/fl^* controls. In *Ntsr1-Cre:tdTomato^fl^:Tle4^fl/fl^* mice, reprogrammed SCPN (tdTomato^+^-Fb^+^ in layers V_ex_ and VI), upregulate CTIP2 (solid arrowheads), and downregulate FOG2 (open arrowheads). (E-F) Visualization of tdTomato^+^ axons from Ntsr1-Cre-expressing neurons in axial sections of the pyramidal tract at P28. Many tdTomato^+^ axons are present in the pyramidal tract of *Cre:tdTomato^fl^:Tle4^fl/fl^,* but not in *Ntsr1-Cre:tdTomato^fl^:Tle4^fl/wt^* controls. Scale bars, 500 μm (a-b, low magnification images), 250 μm (c-d, low magnification), 20 μm (b, d, high magnification), 1 mm (e).

These data indicate that both molecular and projection identity can be “reprogrammed” in CThPN at a relatively late stage when their axons are already innervating the thalamus. To investigate this plasticity of CThPN-to-SCPN identity further, in particular whether axonal projections of reprogrammed SCPN (layer V_ex_ SCPN and layer V SCPN) are transitory or stable, we visualized axons in the pyramidal tract of *Ntsr1-Cre^+^* neurons in P28 *Ntsr1-Cre:tdTomato^fl^*:*Tle4^fl/fl^* mice and *Ntsr1-Cre:tdTomato^fl^:Tle4^flx/wt^* control mice. Quite strikingly, while appropriately there are no tdTomato^+^ axons in the pyramidal tract of *Ntsr1-Cre:tdTomato^fl^:Tle4^flx/wt^* controls, there are tdTomato^+^ axons in the CST of *Ntsr1-Cre:tdTomato^fl^*:*Tle4^fl/fl^* mice (Fig. 6e, f), indicating that reprogrammed SCPN that undergo this plasticity of molecular and projection identity maintain durable projections to subcerebral targets.

Next, we investigated whether this same sort of plasticity– reprogramming of CThPN into SCPN– occurs with loss of *Tle4* function even later, after corticothalamic synapses have already become functional. In somatosensory nuclei, corticothalamic synapses elicit EPSPs by P1 (Golshani et al., 1998). We injected AAV-Cre at P3 in one cortical hemisphere and control AAV in the contralateral cortex of *tdTomato^fl^*:*Tle4^fl/fl^* mice; we confirmed that recombination starts ∼24-48 hours after AAV-Cre injection. Remarkably, *Fezf2* and other SCPN molecular controls are still upregulated, and CThPN molecular controls are still downregulated, at this late stage in the AAV-Cre injected area, but not in the contralateral region (Fig. 7a, Supplementary Fig. 7). These results indicate that loss of *Tle4* function reprograms the molecular identity of CThPN even at this stage of functional corticothalamic circuitry. To investigate whether projection identity is also plastic and can be reprogrammed at this relatively mature stage, we performed AAV-Cre and control AAV injections at P3, retrogradely labeled SCPN at P5, and determined their location in cortex at P12. Quite strikingly, there are non-standard, retrogradely labeled SCPN in layer VI (reprogrammed from CThPN) in the AAV-Cre injected area, but not in the contralateral area (Fig. 7b). These reprogrammed SCPN (originally CThPN) are located at a range of depths in layer VI, but we do not identify a distinct layer V_ex_ as in *Ntsr1-Cre:tdTomato^fl^*:*Tle4^fl/fl^* or *Tle4^KO^ mice*.

**Figure 7.**
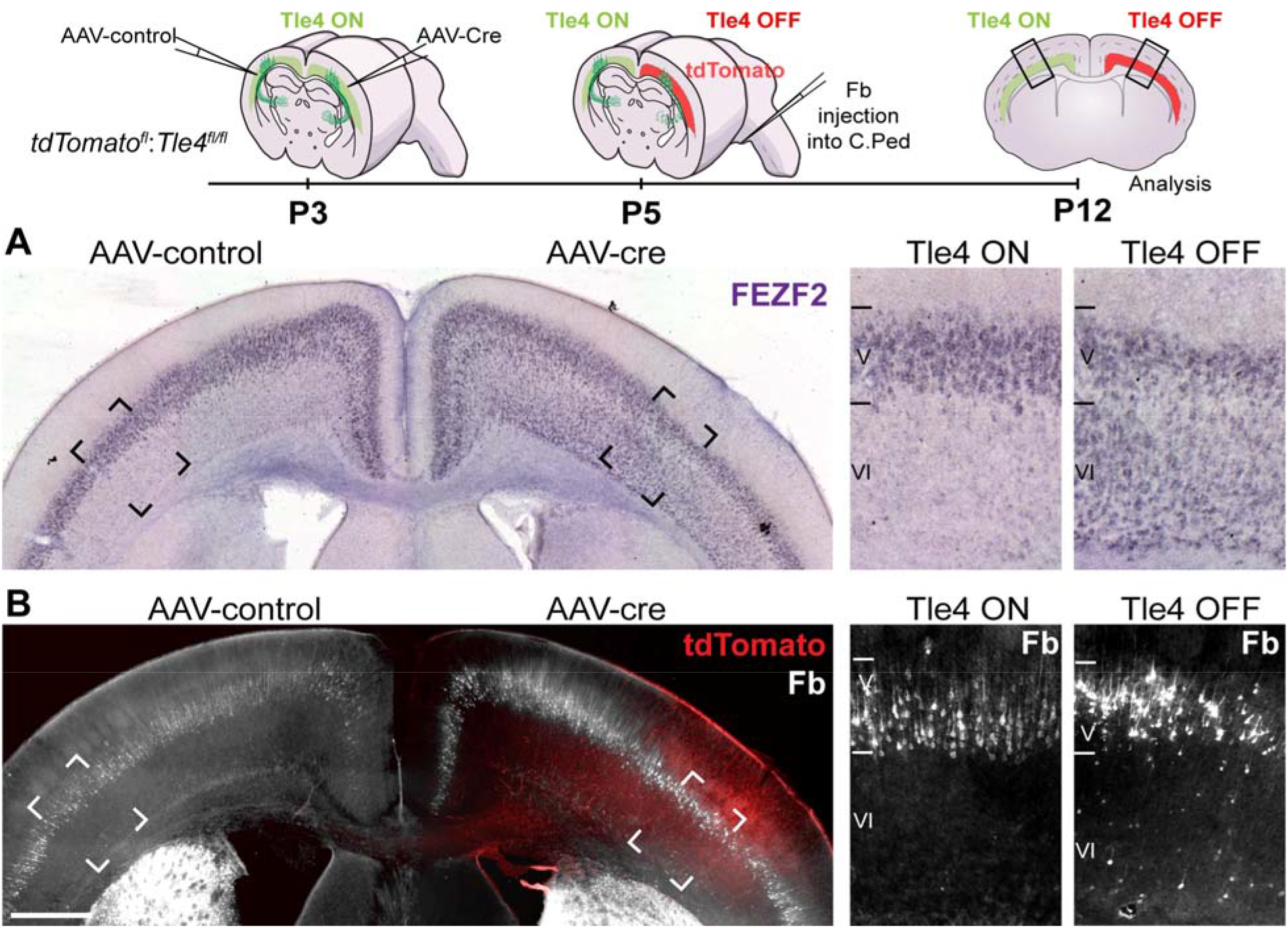
Differentiated CThPN transform their identity in the absence of *Tle4* function during postnatal maturation. CThPN upregulate *Fezf2* and some extend axons into the brainstem after postnatal loss of *Tle4* function during corticothalamic synaptogenesis. Top, schematic of experimental approach. At P3, *tdTomato^fl^:Tle4^fl/fl^* mice are injected with AAV-Cre on one side of the cortex, and AAV-control on the contralateral side. At the time of bilateral injection, *Tle4* is expressed in the areas of viral spread in both hemispheres (*Tle4* ON). At P5, Fb is injected into the cerebral peduncle (C.Ped). At this point, *Tle4* is expressed in the area of viral spread in the hemisphere injected with AAV-control, but not in the contralateral area injected with AAV-Cre (*Tle4* OFF). Analysis of gene expression and retrograde labeling to identify SCPN is performed at P12. (**A**) *Fezf2* ISH demonstrates dramatic upregulation in layer VI of the AAV-Cre injected hemisphere (tdTomato^+^ in b), but not on the contralateral side. (b) Retrograde labeling reveals the presence of Fb^+^ neurons in layer VI (postnatally reprogrammed SCPN) in the AAV-Cre injected area. Higher magnification panels on the right in a and b correspond to the boxed regions in the lower magnification views. Layer V Fb^+^ neurons in both hemispheres appear normal in number and location for SCPN. Scale bar, 500 μm (a-b).

Together, these data indicate that *Tle4* expression is necessary to maintain the molecular and projection identity of CThPN during circuit maturation, and that some CThPN can reprogram both their molecular and projection identity at relatively mature stages of circuit development.

### TLE4 and FEZF2 form a transcription-regulatory complex that modulates activation of *Fezf2*-regulatory regions in developing CThPN

Loss of *Tle4* function, at either embryonic or postnatal stages, results in strong upregulation of *Fezf2* expression by CThPN. Distinct levels of *Fezf2* expression are required for the correct development of distinct corticofugal neuron subtypes– high *Fezf2* expression by SCPN, and low *Fezf2* expression by CThPN (Molyneaux et al., 2005; Chen et al., 2005a; Chen et al., 2005b; Lodato et al., 2014). Abnormally high expression of *Fezf2* by CThPN disrupts their differentiation, and causes some of the phenotypes observed with loss of *Tle4* function. It has been shown that TLE4 and FEZF2 proteins interact directly (Zhang et al., 2014; Tsyporin et al., 2021), and, importantly, that their interaction regulates CThPN development (Tsyporin et al., 2021). However, how TLE4 and FEZF2 proteins function together in regulating CThPN differentiation has not been addressed. We hypothesized that a TLE4-FEZF2 complex might regulate the level of *Fezf2* expression by developing CThPN to achieve the correct *Fezf2* dosage required for CThPN differentiation and distinction from SCPN.

To investigate this hypothesis, we first confirmed that TLE4 and FEZF2 interact in cortical neurons at E15.5 via co-immunoprecipitation (co-IP) of TLE4-Flag and HA-FEZF2 proteins (Fig. 8a). FEZF2 contains an Engrailed1 (Eh1) homology domain, which binds either to the C-terminal WDR domain (also called WD40) or to the N-terminal Q-domain of TLE co-repressors (Buscarlet and Stifani, 2007; Jennings et al., 2006; Chen and Courey, 2000; Shimizu and Hibi, 2009). To determine the specific domain of TLE4 required for interaction with FEZF2, we generated truncated forms of TLE4 lacking either the WDR-domain (Flag-TLE4-ΔWDR) or the Q-oligomerization domain (Flag-TLE4-ΔQ), and investigated their ability to pull-down HA-FEZF2. Co-IP experiments reveal that, while full-length Flag-TLE4 and Flag-TLE4-ΔQ bind and pull down HA-FEZF2, Flag-TLE4-ΔWDR does not pull down HA-FEZF2 (Fig. 8a). These results indicate that the WDR-domain is required for the interaction of TLE4 with FEZF2, and thus for the formation of a putative TLE4-FEZF2 regulatory complex.

**Figure 8.**
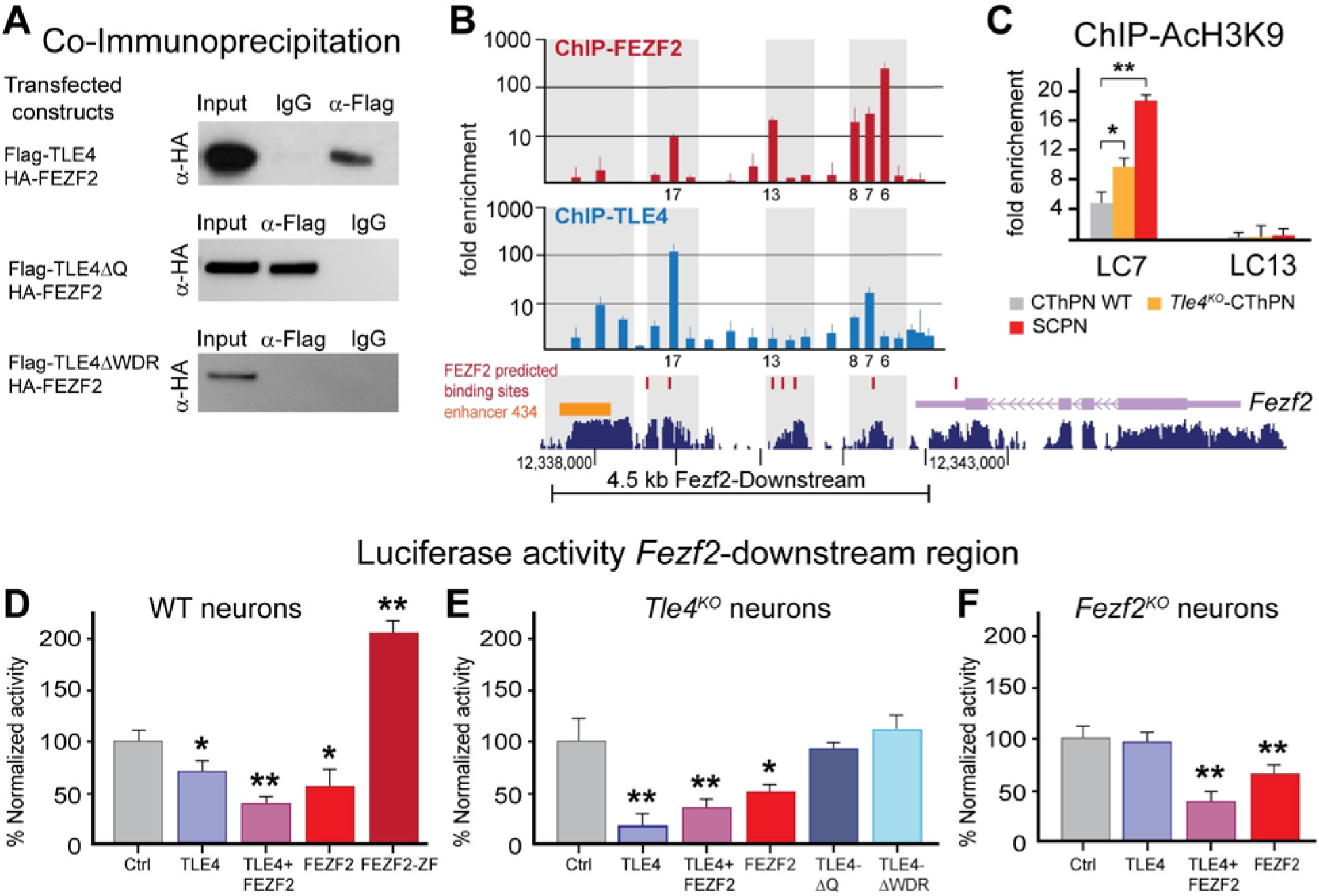
TLE4 and FEZF2 form a regulatory complex that controls *Fezf2* expression level in developing and maturing CThPN. (**A**) Co-IP of protein extracts from E15.5 cortical neurons transfected with *Tle4*-Flag and HA-*Fezf2* constructs reveals binding of TLE4 and FEZF2. HA-FEZF2 protein is detected after pull-downs with full-length TLE4 (Flag-TLE4), and TLE4 lacking the Q-domain (Flag-TLE4ΔQ), but not after pull-down with TLE4 lacking the WDR-domain (Flag-TLE4ΔWDR). Anti-Flag antibody or non-specific IgG were used for pull-downs, and anti-HA antibody for immunoblots. The “Input” lane contains a fraction of the sample before pull-down. (**B**) ChIP-qPCR analysis reveals binding of FEZF2 and TLE4 in the 4.5 kb region downstream of *Fezf2*. Primer sets were designed to span this 4.5 kb downstream region. Amplified loci are numbered from 1 (starting at the *Fezf2* 3’UTR) to 23 (at a location 4.5 kb downstream of *Fezf2*). Loci were aligned to chromosome coordinates. Grey bars map mammalian conserved regions (dark blue, lower gene map) to ChIP-FEZF2 and ChIP-TLE4 enriched loci in the upper two ChIP maps. Predicted *Fezf2* binding sites (red marks in lower map), validated enhancer 434 (orange bar in lower map), and *Fezf2* gene sequence (purple in lower map) are shown for reference. ChIP-FEZF2 and ChIP-TLE4 results are expressed as fold enrichment normalized to control ChIP-IgG for each amplicon. Error bars represent the SEM from 4 biological replicates. (**C**) ChIP for AcH3K9, and qPCR analysis for Locus 7 (LC7) and Locus 13 (LC13). The amount of AcH3K9 in LC7 co-occupied by TLE4 and FEZF2 is significantly higher in both *Tle4^KO^* CThPN and SCPN compared to wildtype CThPN. The amount of AcH3K9 in LC13, not co-occupied by TLE4 and FEZF2, is low and not significantly different between *Tle4^KO^* CThPN, SCPN, and wildtype CThPN. Fold enrichment was determined by normalizing ΔCt values for each locus against the ΔCt values for a constitutive active locus (*Gapdh).* (**D**) Luciferase activity relative to control (cntrl; pGL3-*Fezf2-downstream*-Luciferase alone) in the presence of TLE4, TLE4+FEZF2, FEZF2, and FEZF2-ZF in cortical neurons from E15.5 wildtype embryos. (**E**) Luciferase activity relative to control (pGL3-*Fezf2-downstream*-Luciferase alone) in the presence of TLE4, TLE4+FEZF2, FEZF2, TLE4ΔQ, and TLE4ΔWDR in cortical neurons from E15.5 *Tle4^KO^* embryos. (**F**) Luciferase activity relative to control (pGL3-*Fezf2-downstream*-Luciferase alone) in the presence of TLE4, TLE4+FEZF2, and FEZF2 in cortical neurons from E15.5 *Fezf2^KO^* embryos. Asterisks in **D-F** indicate statistically significant difference compared to control (t-test with Tukey correction for multiple comparison; *p<0.05; **p<0.01).

Next, we investigated whether TLE4 and FEZF2 bind to putative regulatory regions flanking the *Fezf2* locus. Two putative regulatory regions (3 kb-upstream and 4.5 kb-downstream of *Fezf2*) were selected for analysis based on their sequence conservation in mammals, presence of defined enhancers, and presence of predicted FEZF2 binding motifs (Eckler et al., 2014; Sugimoto et al., 2006). To investigate whether TLE4 and FEZF2 bind to loci in these putative *Fezf2* regulatory regions, we performed chromatin immunoprecipitation (ChIP) from E15.5 neurons expressing HA-FEZF2 and Flag-TLE4, followed by qPCR with overlapping primer sets to fully screen the *Fezf2*-downstream region (loci 1-23, Fig. 8b) and the *Fezf2*-upstream region (loci 24-40, not shown). Compared to chromatin pull-down with a non-specific IgG, ChIP with anti-HA antibody (ChIP-FEZF2) was enriched in loci in both the *Fezf2*-upstream and downstream regions. However, ChIP with anti-Flag antibody (ChIP-TLE4) was only enriched in loci downstream of *Fezf2*. Therefore, we focused on the region 4.5 kb-downstream of *Fezf2* as a putative regulatory region potentially controlled by TLE4 and FEZF2.

Loci with more than 10-fold enrichment by both ChIP-FEZF2 and ChIP-TLE4 were considered co-occupied by FEZF2-TLE4, thus potentially regulated by the FEZF2-TLE4 complex. Under this strict selection criterion, we identify two loci co-occupied by FEZF2-TLE4 in the region 4.5 kb downstream of *Fezf2* (locus 7: ChIP-FEZF2 51-fold enrichment over IgG control, ChIP-TLE4 enrichment 29-fold enrichment over IgG control; locus 17: ChIP-FEZF2 10-fold enrichment over IgG control, ChIP-TLE4 153-fold enrichment over IgG control, Fig. 8b).

The presence of two relatively closely neighboring loci co-occupied by FEZF2-TLE4 strongly suggests that the *Fezf2*-downstream region is regulated by the FEZF2-TLE4 complex. However, to directly and rigorously investigate whether the region 4.5 kb downstream of *Fezf2* functions as a distal regulatory element controlled by TLE4 and FEZF2, we performed luciferase reporter assays and tested transcriptional activity from this region under a variety of conditions. We cloned this putative regulatory region into a luciferase reporter construct (pGL3-*Fezf2*-downstream-luc) and transfected E15.5 wildtype neurons with constructs to express either pGL3-*Fezf2*-downstream-luc alone (baseline control), or together with contructs to express TLE4, FEZF2, or TLE4+FEZF2 (Fig. 8d). Relative to baseline activation from *Fezf2*-downstream-luc, either TLE4 or FEZF2 reduces transcriptional activation from this region (Fig. 8d; 28% luc-activity reduction with TLE4, 42% luc-activity reduction with FEZF2, normalized to luc-activity of pGL3-*Fezf2*-downstream-luc). Strikingly, when separate constructs to express both TLE4 and FEZF2 are co-transfected, the reduction is stronger (60% luc-activity reduction with TLE4+FEZF2 compared to baseline). These data reveal that the joint presence of both TLE4 and FEZF2 strongly reduces transcriptional activity from the *Fezf2*-downstream region. Together with our results identifying loci co-occupied by TLE4 and FEZF2 in this region (Fig. 8b) and the known direct interaction between TLE4 and FEZF2 (Fig 8a, and Tsyporin et al., 2021), our data strongly suggest that FEZF2-TLE4 complex regulates transcription in this region.

However, it was theoretically possible that co-expressed TLE4 and FEZF2 might bind to other factors instead of working together to downregulate transcription. To investigate the requirement for complexed TLE4 and FEZF2 for transcriptional regulation in the *Fezf2*-downstream region, we investigated the activity of pGL3-*Fezf2*-downstream-luc in the presence of a truncated form of FEZF2 (FEZF2-ZF), which preserves the zinc-finger DNA-binding domain but lacks the Eh1 domain that binds TLE4. FEZF2-ZF strongly activates transcription from the *Fezf2*-downstream region (Fig. 8d), supporting the interpretation that FEZF2 interaction with TLE4 (or, conceivably, with other co-repressors via the Eh1 domain) is necessary to downregulate expression from this region.

To further investigate the requirement of TLE4 for transcriptional regulation in the *Fezf2*-downstream region, we tested the activity of pGL3-*Fezf2*-downstream-luc in cortical neurons from E15.5 *Tle4^KO^* embryos. The cortical plate at E15.5 is mostly composed of developing deep-layer neurons, which in *Tle4^KO^* embryos strongly express *Fezf2*. Transcription from the *Fezf2*-downstream is strongly downregulated after expression of TLE4 alone or TLE4+FEZF2 in *Tle4^KO^* neurons (81% luc-activity reduction with TLE4 alone, 62% luc-activity reduction with TLE4+FEZF2; Fig. 8e), indicating that the presence of TLE4 is critical for strong repression in this region. Importantly, expression of non-functional forms of TLE4 (TLE4-ΔWDR or TLE4-ΔQ) does not alter transcriptional activity (Fig. 8e). Interestingly, expression of FEZF2 alone in *Tle4^KO^* neurons downregulates the activity of pGL3-*Fezf2*-downstream-luc (49% luc-activity reduction with FEZF2; Fig 8e), suggesting that FEZF2 might recruit other co-repressors, in addition to TLE4, to regulate activity in this region. Further, we investigated the requirement of FEZF2 for regulation of the *Fezf2*-downstream region using neurons from E15.5 *Fezf2^KO^* embryos (Fig. 8f). In *Fezf2^KO^* mice, expression of TLE4 by cortical neurons is reduced (Chen et al., 2005a; Tsyporin et al., 2021). After rescuing expression of FEZF2 in *Fezf2^KO^* neurons, the activity of pGL3-*Fezf2*-downstream-luc is downregulated (33% luc-activity reduction with FEZF2), and, importantly, it is strongly downregulated after co-expression of TLE4+FEZF2 (60% luc-activity reduction with TLE4+FEZF2; Fig 8f). Importantly, transfection of TLE4 alone in *Fezf2^KO^* neurons does not affect the activity of pGL3-*Fezf2*-downstream-luc, indicating that FEZF2 is required for the recruitment of TLE4 as a co-repressor to regulate transcription in the *Fezf2*-downstream region. Collectively, these data indicate that TLE4 and FEZF2 form a regulatory complex that downregulates transcriptional activity from the *Fezf2*-downstream region, and, while FEZF2 conceivably might also recruit other factors, TLE4-mediated repression in this region depends on the presence of FEZF2.

### A TLE4-FEZF2 complex modulates the chromatin state of loci regulating *Fezf2* expression during CThPN differentiation and maturation

The loci enriched in our ChIP-FEZF2 and ChIP-TLE4 experiments likely recruit TLE4-FEZF2 complexes. Since TLE4 is known to bind histone deacetylases (HDACs) to mediate repression (Bandyopadhyay et al., 2014; Brantjes et al., 2001), HDACs recruited to loci co-occupied by FEZF2-TLE4 in the *Fezf2*-downstream regulatory region might likely change the chromatin conformation state at these loci, compacting chromatin and reducing *Fezf2* expression level. Therefore, we investigated whether the acetylation level at co-occupied loci correlates with the level of expression of *Fezf2* by distinct corticofugal subtypes, and with expression of *Fezf2* by CThPN at progressive developmental stages.

Since *Fezf2* is strongly expressed by SCPN and *Tle4^KO^*-CThPN, and is weakly expressed by wildtype CThPN, we compared the acetylation level at a locus co-occupied by TLE4 and FEZF2 (locus 7; Fig. 8b) in these three neuron populations. We isolated wildtype SCPN, CThPN, and *Tle4^KO^*-CThPN from *Rbp4-Cre:tdTomato^fl^*, *Ntsr1-Cre:tdTomato^fl^*, and *Ntsr1-Cre:tdTomato^fl^:Tle4^fl/fl^* P0 pups, respectively, and performed ChIP-qPCR for the open chromatin mark acetylated histone 3-lysine 9 (AcHis3K9). ChIP-AcHis3K9 qPCR values at locus 7 (co-occupied by FEZF2-TLE4) were normalized to ChIP-AcHis3K9 qPCR values at a constitutively active locus (*Gapdh*). In wildtype CThPN, locus 7 is significantly less acetylated than in SCPN (5 fold-enrichment in wildtype CThPN vs. 18 fold-enrichment in SCPN, p<0.01; Fig. 8c). This is consistent with the presence of a FEZF2-TLE4 complex recruiting HDACs to this locus in CThPN, but not in SCPN, in which the lack of TLE4 expression prevents the formation of the complex. Importantly, locus 7 is also significantly more acetylated in *Tle4^KO^*-CThPN than in wildtype CThPN (5 fold-enrichment in wildtype CThPN vs. 9.6 fold-enrichment in *Tle4^KO^*-CThPN; p<0.05; Fig 8C). In important comparison, the acetylation level at a locus of the *Fezf2*-downstream region not co-occupied by TLE4 and FEZF2 (locus 13) is not significantly different between wildtype CThPN, SCPN, and *Tle4^KO^*-CThPN (0.5 fold-enrichment in SCPN, 0.3 fold-enrichment in CThPN, 0.22 fold-enrichment in *Tle4^KO^*-CThPN; Fig. 8c). Together, our data identify that at least one FEZF2-TLE4 co-occupied locus (locus 7) is differentially acetylated in CThPN, SCPN, and *Tle4^KO^*-CThPN, and that reduced acetylation at this locus correlates with lower *Fezf2* expression in these corticofugal neuron subtypes. These results strongly support the hypothesis that FEZF2-TLE4 complexes recruit HDACs to *Fezf2* regulatory loci, reducing their activation and *Fezf2* expression.

Next, we investigated whether the acetylation level at locus 7 changes during CThPN embryonic and postnatal differentiation and maturation, reflecting *Fezf2* expression changes by CThPN during this period. Developing embryonic CThPN express *Fezf2* strongly, after which *Fezf2* is progressively reduced postnatally and maintained at a low level (Supplementary Fig. 8a). To investigate the acetylation level at the FEZF2-TLE4 co-occupied locus 7 in developing CThPN, we isolated E15.5 cortical neurons, P0 CThPN, and P6 CThPN (from P0 and P6 *Ntsr1-Cre:tdTomato^fl^* mice), and performed ChIP-AcHis3K9 followed by qPCR for this locus. Strikingly, acetylation at locus 7 drastically decreases postnatally compared to the embryonic stage (152 fold-enrichment at E15.5, 58 fold-enrichment at P0, 13.8 fold-enrichment at P6, fold-enrichment values normalized to *Gapdh*; p<0.01; Supplementary Fig. 8b). Importantly, the acetylation level of a nearby locus not co-occupied by FEZF2-TLE4 (locus 13) does not change over time. These data indicate that progressive deacetylation at loci regulated by a FEZF2-TLE4 complex correlates with the progressive reduction of *Fezf2* expression by CThPN during differentiation.

Collectively, these studies reveal a powerful epigenetic mechanism mediated by FEZF2-TLE4 complexes that regulates *Fezf2* expression to stage-appropriate levels during CThPN differentiation and maturation. This mechanism contributes importantly to establishing and maintaining the appropriate levels of *Fezf2* expression necessary for the distinction of CThPN and SCPN subtypes, and to maintaining the low level of *Fezf2* expression necessary for postnatal CThPN maturation, maintenance, and stability.

## Discussion

Our results reveal both that *Tle4* is necessary during embryonic development for the acquisition of precise CThPN identity by early-born cortical neurons, and that *Tle4* is necessary for differentiated CThPN to maintain their distinct identity during postnatal maturation. We further identify mechanistically that TLE4 is recruited by FEZF2 to control chromatin conformation at specific loci regulating the expression level of *Fezf2* in CThPN during embryonic and postnatal development. This epigenetic mechanism contributes importantly to the distinction of CThPN and SCPN subtypes, and ensures appropriate maturation of CThPN, thus neuronal diversity of connectivity, circuitry, and function.

Consistent with recent observations in *Tle4^LacZ/LacZ^* mice (Tsyporin et al., 2021) (*LacZ* insertion into intron 4, disrupting expression), our experiments reveal substantial abnormalities of CThPN development in *Tle4^KO^* mice (deletion after exon 1, constitutive null, no functional protein) (Wheat et al., 2014). In both *Tle4* mutant mouse models, the expression of important SCPN molecular controls are upregulated by CThPN, the expression of CThPN molecular controls are downregulated, and the dendritic development of CThPN is abnormal. Also in both models, projection to the thalamus appears grossly normal, indicating that *Tle4* does not directly instruct CThPN axons to extend into the thalamus.

Extending these results to the level of subtype diversity, in *Tle4^KO^* mice, specific subsets of neurons fate-switch and become SCPN. These neurons not only exhibit abnormal gene expression, but also abnormal SCPN-like somatic morphology, with axonal projections to the brainstem and/or spinal cord. While some phenotypic differences between mutant models are expected due to differing gene-targeting approaches, overall, the results of loss of *Tle4* function across both mutant models largely overlap, revealing the central importance of *Tle4* function in CThPN development.

In the absence of *Tle4* function, strong upregulation of *Fezf2* by developing CThPN contributes importantly to their abnormal differentiation. *Fezf2* is critical for differentiation of corticofugal neurons, in particular to instruct SCPN identity (Molyneaux et al., 2005; Chen et al., 2005a; Chen et al., 2005b). Elegant experiments by Tsyporin et al., 2021 find that expression of a repressive chimeric form of Fezf2 (*Fezf2-EnR,* consisting of the Fezf2 DNA-binding domain fused to the repressive domain of engrailed protein) in *Tle4^LacZ/LacZ^* mutants restored the expression level of important SCPN controls upregulated by CThPN, and the dendritic and functional abnormalities of CThPN in *Tle4^LacZ/LacZ^* mutants. These experiments suggested that FEZF2 and TLE4 together regulate the molecular, morphological, and functional differentiation of CThPN, and highlighted the importance of *Fezf2* expression in regulating these multiple aspects of CThPN development (Tsyporin et al., 2021).

Our work now mechanistically identifies that FEZF2 and TLE4 function together to control CThPN differentiation by regulating the level of expression of *Fezf2* by CThPN. Our work both reinforces the results of the recent study from Tsyporin et al., 2021, and substantially extends them mechanistically to reveal the embryonic and continued postnatal collaborative functions of FEZF2 and TLE4 in CThPN differentiation, in particular in the critical regulation of FEZF2 dosage for appropriate corticofugal neuron distinction and maturation.

TLE4 displays multiple functions in CThPN development, including regulating some gene expression independently of FEZF2. Interestingly, in *Tle4^LacZ/LacZ^*; *Fezf2-EnR* rescue\ experiments (Tsyporin et al., 2021), the expression of important CThPN controls such as *Fog2* and *Foxp2* was not restored. The strong upregulation of *Fezf2* by CThPN in the absence of *Tle4* function likely contributes to the fate-conversion of some CThPN into SCPN observed in *Tle4^KO^* mice, in addition to underlying some of the abnormal phenotypes in both *Tle4* mutants. Intriguingly, *Fezf2* is strongly upregulated by virtually all developing CThPN in *Tle4^KO^* cortex, including those that do not fate-convert to become SCPN. Therefore, in addition to *Fezf2*, other important controls dysregulated in the absence of *Tle4* function likely contribute to CThPN plasticity and their capacity to fate-switch to become SCPN.

The data reported here support the emerging interpretation that intrinsic molecular diversity of CThPN contributes to their differential plasticity, rendering some specific CThPN subsets more amenable to fate-switch into SCPN (Galazo et al., 2016; Tomassy et al., 2010). The result that fate-converted SCPN in layer ‘V_ex_’ are more similar to normal wildtype SCPN than fate-converted SCPN in layer VI is consistent with this idea that distinct CThPN subsets have intrinsic gene expression differences that importantly contribute to their differential plasticity, and that these transcriptional differences might be critical for correct implementation of the SCPN identity program following upregulation of *Fezf2*.

In striking contrast to prior work with other regulators of CThPN identity, our findings in *Tle4^KO^* mice reveal that there are fate-converted SCPN with the molecular and connectivity characteristics of normal SCPN, including complex area-specific axonal projection patterns. Acquisition of abnormal SCPN identity by neurons that normally become CThPN has been described in mutant mice with targeted gene deletion of *Tbr1, Sox5,* or *Couptf-1*. However, in these mutants, subcerebral projections are abnormal, extending to incorrect targets or being unstable and pruned over time (Hevner et al., 2001; Mckenna et al., 2011; Han et al., 2011; Lai et al., 2008; Kwan et al., 2008; Tomassy et al., 2010). Perhaps the broad *Tle4* function reported here contributes to more broadly encompassing regulation of CThPN vs. SCPN identity, differentiation, connectivity, and likely circuit function.

*Tle4* function and its mechanisms of both embryonic and postnatal transcriptional regulation are important beyond corticofugal differentiation, potentially enabling control over directed differentiation or trans-differentiation of corticofugal lineages. There is substantial interest in understanding how to differentiate specific neuron subtypes *in vitro* that faithfully reflect the characteristics of the neuron subtypes *in vivo* (Rouaux et al., 2012; Barker et al., 2018). There is also substantial interest in altering or reprogramming existing neuron types into others to replace or compensate for those that are selectively vulnerable in disease, e.g. SCPN in amyotrophic lateral sclerosis (ALS) / motor neuron disease (MND). For such hypothetical approaches to offer optimal benefit, conversion to, e.g. SCPN, would need to reflect broad connectivity and molecular features of the desired new neuron subtype, not just a few molecular markers. The regulation by *Tle4* loss-of-function of broad SCPN molecular expression and connectivity offers promise in this regard. Some lines of evidence indicate that at least some aspects of neuron subtype identity remain plastic, and can be altered for at least a short time after initial development. Some aspects of cortical neuron identity, including gene expression, dendritic morphology, and electrophysiological properties have been reprogrammed early-postnatally in mice (Rouaux and Arlotta, 2013; De La Rossa et al., 2013). However, only embryonically has projection identity of cortical projection neurons been successfully reprogrammed. Mis-expression of *Fezf2* in superficial layer CPN at e17.5 induces expression of some corticofugal genes and extension of axonal projections to subcortical structures (Rouaux and Arlotta, 2013). However, it is important to note that, at this stage, CPN are immature and are still extending axons dorsal to the striatum toward the midline, amenable to re-direction ventrally into the striatum. Re-direction of SCPN connectivity by cortical neurons has not been reported at later developmental or postnatal stages.

Our experiments manipulating neuronal identity by inducing *Tle4* null mutation postnatally reveal a substantially more extended time window for neuron plasticity and projection identity reprogramming in CThPN compared to other cortical neuron subtypes. It is striking that some CThPN can seemingly fully change their axonal connectivity even after P3, especially since they are the earliest-born cortical neuron subtype, and they are already establishing synaptic connections in the thalamus at this postnatal stage. It will be interesting in the future to understand whether this extended time window for identity reprogramming exhibited by CThPN reflects intrinsically enhanced capacity for plasticity of this specific subtype, or perhaps reflects that identity reprogramming might be more effectively accomplished between neuron subtypes with similar developmental trajectories (in this case between the corticofugal subtypes CThPN and SCPN), and/or might extend for a longer period of time between closely related subtypes.

The epigenetic mechanism identified here by which TLE4 mediates deacetylation at specific loci regulating *Fezf2* expression contributes to establishing the initial sharp distinction between the CThPN and SCPN corticofugal subtypes, and to maintaining CThPN identity during postnatal maturation. Epigenetic regulation is known to progressively restrict cell fate potential during development (Ziller et al., 2015). Early in cortical development, progenitors and postmitotic deep-layer corticofugal neurons strongly express FEZF2. Developing CThPN express TLE4 as soon as they are positioned in the cortical plate. The TLE4-FEZF2 complex identified here likely starts functioning at this early stage to progressively reduce FEZF2 expression level and thus direct correct acquisition of CThPN identity. Postnatally, continuous *Tle4* expression and function of this TLE4-FEZF2 complex in CThPN ensure low-level expression of *Fezf2*, which our experiments identify as necessary to maintain CThPN identity and avoid reprogramming into SCPN. In mice, maintenance of neuron subtype identity characteristics has been studied primarily in the dopaminergic and serotonergic systems, in which it is found that transcription factors controlling maintenance of neurotransmitter identity are also crucial for neurotransmitter identity specification (Deneris and Hobert, 2014). Here, we identify both that the TLE4-FEZF2 complex is similarly required for acquisition and maintenance of CThPN identity, and that the TLE4-FEZF2 complex controls broad molecular and connectivity characteristics of CThPN vs. SCPN subtype identity, rather than only neurotransmitter expression.

It is intriguing to note that the epigenetic mechanism identified here for transcriptional regulation of Fezf2 expression functions as a negative feedback loop, by which FEZF2 recruits TLE4 as co-repressor for autorepression. In invertebrates, direct positive feedback autoregulation has been described as a mechanism to ensure long-term expression of genes involved in neuron identity maintenance (Deneris and Hobert, 2014; Ptashne, 2014; Etchberger et al., 2007; Wenick and Hobert, 2004). It appears now that similar autoregulation feedback mechanisms function centrally in neuron subtype identity development, maintenance, and potentially plasticity in the mammalian cortex, though perhaps these mechanisms are more complex in cortex. The presence of transcriptional co-regulators such as TLE4 could provide a mechanism to dynamically control the outcome of autoregulation feedback loops in distinct cellular contexts, e.g. between positive feedback and negative feedback to increase or decrease expression. This could theoretically enable more precise control of expression levels of critical genes required to regulate neuron subtype identity and diversity in the cerebral cortex.

In addition to the central function of *Tle4* in acquisition of CThPN identity by early-born cortical neurons during embryonic development, our work reveals a novel and critical function of *Tle4* in controlling CThPN identity, plasticity, and stability during postnatal maturation. Continuous *Tle4* expression is required to ensure function of the TLE4-FEZ2 complex to actively maintain CThPN identity. Similarly, *Tbr1* is critical for development of CThPN identity by early-born cortical neurons, and its continued expression is important for maintaining CThPN molecular identity, dendritic morphology, and synaptic properties (Fazel Darbandi et al., 2018). Together, *Tle4* and *Tbr1* might regulate complementary mechanisms controlling maintenance of broad molecular and input-output connectivity characteristics of CThPN identity. Interestingly, other important developmental controls over cortical neuron subtype identity, including *Fezf2* (Molyneaux et al., 2005; Chen et al., 2005a; Chen et al., 2005b), *Ctip2* (Arlotta et al., 2005), *Sox5* (Lai et al., 2008; Kwan et al., 2008), *Couptf1* (Tomassy et al., 2010), and *Fog2* (Galazo et al., 2016), are also expressed continuously during postnatal cortical development (<u>mouse.brain-map.org</u>), so perhaps they might also have additional functions during neuron maturation and maintenance that have not been investigated yet. Our work here revealing multiple functions of *Tle4* over an extended temporal course underscores the importance of regulating not only early neuron identity acquisition, but also neuron subtype identity stability during maturation for correct long-term functions of neurons in complex circuits.

In summary, the experiments we report here elucidate two critical and separable functions of *Tle4* in cortical development and maintenance of correct circuitry. Early in corticogenesis, *Tle4* is required to promote acquisition of corticothalamic molecular and cellular identity, centrally including connectivity, and block emergence of subcerebral / corticospinal projection neuron identity. Postnatally, *Tle4* is required to maintain corticothalamic molecular identity and proper connectivity during circuit maturation. We further identify a powerful epigenetic mechanism by which TLE4 controls the activation state of loci regulating the level of *Fezf2* expression by corticothalamic neurons during embryonic and postnatal development and maintenance. This novel TLE4-FEZF2 negative feedback autorepression mechanism contributes importantly to the developmental distinction of cortical output (corticofugal) subtypes, and ensures appropriate maturation and maintenance of CThPN with their functionally critical connectivity and circuitry.

## Supporting information

Supplemental Material_Galazo, Sweetser, Macklis

## Author Contributions

M.J.G. and J.D.M. conceived the overall project and experiments; D.A.S. generated *Tle4* mouse lines; M.J.G. designed and performed the experiments; M.J.G. analyzed the data; M.J.G. and J.D.M. interpreted the data, integrated the findings, and wrote the manuscript. All authors read and approved the final manuscript.

## Acknowledgments

We thank E. L. Gorstein, C. Greppi, I. Florea, and K. Yee for superb technical assistance; members of the Macklis laboratory for scientific discussions and helpful suggestions; the Harvard Center for Biological Imaging for infrastructure and support; S.K. McConnell for generous sharing of *Fezf2-PLAP* mice. This work was supported by grants from the National Institutes of Health (R01 NS045523 with additional infrastructure supported by NS075672, NS049553, NS104055, and DP1 NS106665) to J.D.M.

M. J. G. was partially supported by fellowships from CajaMadrid Foundation and the Spanish Ministry of Education, Brain and Behavior Foundation, Tulane COR research program, and Louisiana Board of Regents (LEQSF(2021-24)-RD-A-14). J.D.M. was also supported as an Allen Distinguished Investigator of the Paul G. Allen Frontiers Group.

## Experimental Procedures

### Mice

All mouse studies were approved by the Harvard University and Massachusetts General Hospital IACUCs, and were performed in accordance with institutional and federal guidelines. *Tle4*^KO^ and *Tle4^floxed^* mice were described in (Wheat et al., 2014). *Emx1*-Cre (stock number 005628) and tdTomato*^fl^* (Ai14*-Rosa26-tdTomato^fl-stop-fl^*; stock number 007914) mice were purchased from Jackson Laboratories. *Ntsr1*-Cre (RRID:MGI_3836636, stock number 030648-UCD) and *Rbp4*-Cre ((RRID:MGI_4367067, stock number 031125-UCD) mice were generated by the GENSAT project (Gong et al., 2007), and were purchased from MMRRC. *Fezf2-PLAP* mice were the generous gift of Dr. S. McConnell at Stanford University (Chen et al., 2005a). The morning of vaginal plug detection was designated E0.5, and the day of birth as P0. Unless noted otherwise, all experiments with *Tle4*^KO^ were controlled with *Tle4^+/+^,* although no abnormal cortical phenotypes were observed in *Tle4^+/−^*.

### Immunocytochemistry and *in situ* hybridization

Brains were fixed and stained using standard methods (Molyneaux et al., 2005; Lai et al., 2008). Primary antibodies and dilutions used: rat anti-BrdU, 1:750 (Accurate Chemical and Scientific Corporation Cat# OBT-0030 RRID:AB_2341179); rat anti-CTIP2, 1:500 (Abcam Cat# ab28448, RRID:AB_1140055); rabbit anti-DARPP-32, 1:250 (Cell Signaling Technology Cat# 2306S, RRID:AB_823479); rabbit anti-FOG2, 1:250 (Santa Cruz Cat# sc-10755, RRID:AB_2218978); goat anti-NURR1, 1:100 (R&D Systems Cat# AF2156, RRID:AB_2153894); rabbit anti-TBR1, 1:500 (Abcam Cat# ab31940, RRID:AB_2200219); rabbit anti-FOXP2 1:1000 (Abcam Cat# ab1307, RRID:AB_1268914); rabbit anti-TLE4, 1:200 (Santa Cruz Biotechnology Cat# sc-9125, RRID:AB_793141). Rat anti-BrdU requires antigen retrieval, as described in (Magavi and Macklis, 2008). *In situ* hybridization was performed as previously described in (Arlotta et al., 2005). cDNA clones for riboprobes are listed in (Galazo et al., 2016).

### Retrograde Labeling

Projection neurons were retrograde labeled from their axon terminals by pressure injection of Alexa-conjugated Cholera Toxin B (Ctb, Invitrogen) or Fast Blue (Polysciences) at distinct stages. In postnatal pups, injections into either the cerebral peduncle, pyramidal tract, cervical spinal cord, dorsal thalamus, or cortex were performed under ultrasound backscatter microscopy guidance (Vevo 770, VisualSonics, Toronto) at P3 or P5, depending on the experimental condition. In adults, stereotaxic injections into the superior colliculus or pyramidal tract-pontine nuclei were performed following coordinates from (Paxinos, 2001) Paxinos and Franklin, *The Mouse Brain in Stereotaxic Coordinates, 2^nd^ edition*, 2001. For retrograde labeling experiments, n=4 to 6 for each genotype.

### BrdU birthdating

Timed pregnant females were injected with bromodeoxyuridine (50 mg/kg, i.p) at E11.5, E12.5, E13.5, or E14.5. For experiments in which BrdU birthdating was combined with SCPN retrograde labeling, Ctb-Alexa555 was injected into the cerebral peduncle at P3. Brains were collected at P6, and processed for either 1) BrdU and CTIP2 or 2) BrdU and FOG2 dual immunocytochemistry. Co-localization and quantification of SCPN, BrdU^+^, and CTIP2^+^ or FOG2^+^ neurons, and their distribution within cortical layers was analyzed using established methods (Molyneaux et al., 2005; Lai et al., 2008; Woodworth et al., 2016). For BrdU-labeling experiments, n=4 to 6 for each genotype.

### Viral injections

Viral injections were performed under ultrasound backscatter microscopy guidance (Vevo 770, VisualSonics, Toronto) at P3. *TdTomato^fl^:Tle4^fl/^*^fl^ mice were injected with 70 μl of AAV-Cre, distributed in several injection sites in one hemisphere of the cortex, and 70 μl of AAV-control in the contralateral hemisphere. AAV-Cre or AAV-control serotype 2.1 were obtained from the Massachusetts General Hospital Viral Core.

### Protein Co-Immunoprecipitation

Dissociated E15.5 cortical neurons were co-transfected with *HA-Fezf2* and either *Flag-Tle4*, *Flag-Tle4ΔQ, or Flag-Tle4ΔWDR* constructs, using Amaxa nucleofection (Lonza). Protein IP was performed using Protein-A/G agarose beads (Pierce), following standard methods (Galazo et al., 2016; Huggins et al., 2001). Mouse anti-Flag and non-specific mouse IgG (4mg/ml; Sigma F7425; RRIB:AB_439687) were used for pull-down. Rabbit anti-HA (1:2000, Covance, MMS-101R, RRID:AB_291262,) was used for immunoblotting.

### ChIP-qPCR

For ChIP-FEZF2 and ChIP-TLE4, dissociated E15.5 cortical neurons were co-transfected with *HA-Fezf2* and *Flag-Tle4* constructs, using Amaxa nucleofection (Lonza). ChIP-qPCR was performed following standard methods. In brief, double crosslinking (protein-protein, and protein-DNA) was performed using EGS cross-linker (ethylene glycol bis-sulfosuccinimidyl succinate) or formaldehyde, respectively, according to (Forster et al., 2014). Chromatin shearing was optimized for the amplicon size. Anti-Flag (Sigma F7425; RRIB:AB_439687), anti-HA (Covance, MMS-101R, RRID:AB_291262), or non-specific IgG antibodies were used for ChIP bound to protein G/A beads. Pulled-down chromatin was purified and quantified by qPCR (LightCycler Fast start DNA Master SYBER Green I, Roche) with overlapping sets of primers designed to cover a region approximately 4.5 kb downstream of Fezf2 (chr14: 12,337,560..12,342,460). Amplicon average size is 300 bp. A list of chromosome coordinates for each amplicon is provided in Supplementary Table 1. For each amplicon, the fold enrichment listed is of anti-Flag pull-down over IgG or anti-HA pull-down over IgG, using the ΔCt method.

ChIP-AcH3K9 was performed using either wildtype cortical neurons obtained from E15.5 embryos, P0, or P3 pups, or using FACS-purified CThPN, SCPN, or *Tle4^KO^*-CThPN obtained from *Ntsr1-Cre:tdTomato^fl^*, *Rbp4-Cre:tdTomato^fl^*, or *Ntsr1-Cre:tdTomato^fl^:Tle4^KO^* E15.5 embryos, respectively. FACS purification was performed as described in Arlotta et al., 2005^22^. Anti-AcH3K9 (Millipore 17-658, RRID:AB_1587124) or non-specific IgG antibodies were used for ChIP bound to protein G/A beads. Pulled-down chromatin was quantified by qPCR. Fold enrichment was determined by normalizing ΔCt values for each locus against IgG control, and against a constitutive active locus (*Gapdh*). For ChIP experiments, n = 4 independent replicates were used per group.

### Luciferase Assay

The 4.5 kb downstream of *Fezf2* was obtained from a BAC clone (RP23-141E17), and cloned into pGL3-Firefly luciferase upstream of the SV40 promoter (pGL3-luc; Promega) using a Gibson assembly kit (New England BioLabs) to generate pGL3-*Fezf2*-downstram-luc. Dissociated neurons from dissected E15.5 wildtype, *Tle4^KO^*, or *Fezf2^KO^* cortices were nucleofected with pGL3-*Fezf2*-downstram-luc, Renilla luciferase vector, and either *Tle4, Fezf2, Fezf2-ZF, Tle4ΔQ, or Tle4ΔWDR* expression plasmids, alone or in combination, depending on the experimental condition (5 to 7 independent biological replicates per condition). Luciferase activities were assayed 48 h later using the Dual-Glo system (Promega). Firefly/Renilla luciferase ratio was calculated for each condition and referred to as the percentage relative to the baseline activity of pGL3-*Fezf2*-downstram-luc.

### Statistics

Sample sizes were similar to those used in previous publications from our group and others, so no additional statistics were used to determine group sample sizes. Two-tailed t-student, or ANOVA with Tukey correction for pairwise comparisons were used. Values are represented as means ± SEM. Asterisks indicate significance (*p<0.05, **p<0.001)

## References

Allen, T. & Lobe, C. G. 1999. A comparison of Notch, Hes and Grg expression during murine embryonic and post-natal development. Cell Mol Biol (Noisy-le-grand), 45, 687–708.

Arlotta, P., Molyneaux, B. J., Chen, J., Inoue, J., Kominami, R. & Macklis, J. D. 2005. Neuronal subtype-specific genes that control corticospinal motor neuron development in vivo. Neuron, 45, 207–21.

Bandyopadhyay, S., Valdor, R. & Macian, F. 2014. Tle4 regulates epigenetic silencing of gamma interferon expression during effector T helper cell tolerance. Mol Cell Biol, 34, 233–45.

Barker, R. A., Gotz, M. & Parmar, M. 2018. New approaches for brain repair-from rescue to reprogramming. Nature, 557, 329–334.

Brantjes, H., Roose, J., Van De Wetering, M. & Clevers, H. 2001. All Tcf HMG box transcription factors interact with Groucho-related co-repressors. Nucleic Acids Res, 29, 1410–9.

Buscarlet, M. & Stifani, S. 2007. The ‘Marx’ of Groucho on development and disease. Trends Cell Biol, 17, 353–61.

Chen, B., Schaevitz, L. R. & Mcconnell, S. K. 2005a. Fezl regulates the differentiation and axon targeting of layer 5 subcortical projection neurons in cerebral cortex. Proc Natl Acad Sci U S A, 102, 17184–9.

Chen, G. & Courey, A. J. 2000. Groucho/TLE family proteins and transcriptional repression. Gene, 249, 1–16.

Chen, J. G., Rasin, M. R., Kwan, K. Y. & Sestan, N. 2005b. Zfp312 is required for subcortical axonal projections and dendritic morphology of deep-layer pyramidal neurons of the cerebral cortex. Proc Natl Acad Sci U S A, 102, 17792–7.

Chevee, M., Robertson, J. J., Cannon, G. H., Brown, S. P. & Goff, L. A. 2018. Variation in Activity State, Axonal Projection, and Position Define the Transcriptional Identity of Individual Neocortical Projection Neurons. Cell Rep, 22, 441–455.

De La Rossa, A., Bellone, C., Golding, B., Vitali, I., Moss, J., Toni, N., Luscher, C. & Jabaudon, D. 2013. In vivo reprogramming of circuit connectivity in postmitotic neocortical neurons. Nat Neurosci, 16, 193–200.

Deneris, E. S. & Hobert, O. 2014. Maintenance of postmitotic neuronal cell identity. Nat Neurosci, 17, 899–907.

Eckler, M. J., Larkin, K. A., Mckenna, W. L., Katzman, S., Guo, C., Roque, R., Visel, A., Rubenstein, J. L. & Chen, B. 2014. Multiple conserved regulatory domains promote Fezf2 expression in the developing cerebral cortex. Neural Dev, 9, 6.

Etchberger, J. F., Lorch, A., Sleumer, M. C., Zapf, R., Jones, S. J., Marra, M. A., Holt, R. A., Moerman, D. G. & Hobert, O. 2007. The molecular signature and cis-regulatory architecture of a C. elegans gustatory neuron. Genes Dev, 21, 1653–74.

Fazel Darbandi, S., Robinson Schwartz, S. E., Qi, Q., Catta-Preta, R., Pai, E. L., Mandell, J. D., Everitt, A., Rubin, A., Krasnoff, R. A., Katzman, S., Tastad, D., Nord, A. S., Willsey, A. J., Chen, B., State, M. W., Sohal, V. S. & Rubenstein, J. L. R. 2018. Neonatal Tbr1 Dosage Controls Cortical Layer 6 Connectivity. Neuron, 100, 831–845 e7.

Forster, N., Saladi, S. V., Van Bragt, M., Sfondouris, M. E., Jones, F. E., Li, Z. & Ellisen, L. W. 2014. Basal cell signaling by p63 controls luminal progenitor function and lactation via NRG1. Dev Cell, 28, 147–60.

Frantz, G. D. & Mcconnell, S. K. 1996. Restriction of late cerebral cortical progenitors to an upper-layer fate. Neuron, 17, 55–61.

Galazo, M. J., Emsley, J. G. & Macklis, J. D. 2016. Corticothalamic Projection Neuron Development beyond Subtype Specification: Fog2 and Intersectional Controls Regulate Intraclass Neuronal Diversity. Neuron, 91, 90–106.

Golshani, P., Warren, R. A. & Jones, E. G. 1998. Progression of change in NMDA, non-NMDA, and metabotropic glutamate receptor function at the developing corticothalamic synapse. J Neurophysiol, 80, 143–54.

Gong, S., Doughty, M., Harbaugh, C. R., Cummins, A., Hatten, M. E., Heintz, N. & Gerfen, C. R. 2007. Targeting Cre recombinase to specific neuron populations with bacterial artificial chromosome constructs. J Neurosci, 27, 9817–23.

Grant, E., Hoerder-Suabedissen, A. & Molnar, Z. 2012. Development of the corticothalamic projections. Front Neurosci, 6, 53.

Greig, L. C., Woodworth, M. B., Galazo, M. J., Padmanabhan, H. & Macklis, J. D. 2013. Molecular logic of neocortical projection neuron specification, development and diversity. Nat Rev Neurosci, 14, 755–69.

Guillery, R. W. & Sherman, S. M. 2002. Thalamic relay functions and their role in corticocortical communication: generalizations from the visual system. Neuron, 33, 163–75.

Han, W., Kwan, K. Y., Shim, S., Lam, M. M., Shin, Y., Xu, X., Zhu, Y., Li, M. & Sestan, N. 2011. TBR1 directly represses Fezf2 to control the laminar origin and development of the corticospinal tract. Proc Natl Acad Sci U S A, 108, 3041–6.

Hevner, R. F., Shi, L., Justice, N., Hsueh, Y., Sheng, M., Smiga, S., Bulfone, A., Goffinet, A. M., Campagnoni, A. T. & Rubenstein, J. L. 2001. Tbr1 regulates differentiation of the preplate and layer 6. Neuron, 29, 353–66.

Huggins, G. S., Bacani, C. J., Boltax, J., Aikawa, R. & Leiden, J. M. 2001. Friend of GATA 2 physically interacts with chicken ovalbumin upstream promoter-TF2 (COUP-TF2) and COUP-TF3 and represses COUP-TF2-dependent activation of the atrial natriuretic factor promoter. J Biol Chem, 276, 28029–36.

Jacobs, E. C., Campagnoni, C., Kampf, K., Reyes, S. D., Kalra, V., Handley, V., Xie, Y. Y., Hong-Hu, Y., Spreur, V., Fisher, R. S. & Campagnoni, A. T. 2007. Visualization of corticofugal projections during early cortical development in a tau-GFP-transgenic mouse. Eur J Neurosci, 25, 17–30.

Jennings, B. H., Pickles, L. M., Wainwright, S. M., Roe, S. M., Pearl, L. H. & Ish-Horowicz, D. 2006. Molecular recognition of transcriptional repressor motifs by the WD domain of the Groucho/TLE corepressor. Mol Cell, 22, 645–55.

Joshi, P. S., Molyneaux, B. J., Feng, L., Xie, X., Macklis, J. D. & Gan, L. 2008. Bhlhb5 regulates the postmitotic acquisition of area identities in layers II-V of the developing neocortex. Neuron, 60, 258–72.

Kwan, K. Y., Lam, M. M., Krsnik, Z., Kawasawa, Y. I., Lefebvre, V. & Sestan, N. 2008. SOX5 postmitotically regulates migration, postmigratory differentiation, and projections of subplate and deep-layer neocortical neurons. Proc Natl Acad Sci U S A, 105, 16021–6.

Lai, T., Jabaudon, D., Molyneaux, B. J., Azim, E., Arlotta, P., Menezes, J. R. & Macklis, J. D. 2008. SOX5 controls the sequential generation of distinct corticofugal neuron subtypes. Neuron, 57, 232–47.

Liu, X. B., Murray, K. D. & Jones, E. G. 2004. Switching of NMDA receptor 2A and 2B subunits at thalamic and cortical synapses during early postnatal development. J Neurosci, 24, 8885–95.

Lodato, S. & Arlotta, P. 2015. Generating neuronal diversity in the mammalian cerebral cortex. Annu Rev Cell Dev Biol, 31, 699–720.

Lodato, S., Molyneaux, B. J., Zuccaro, E., Goff, L. A., Chen, H. H., Yuan, W., Meleski, A., Takahashi, E., Mahony, S., Rinn, J. L., Gifford, D. K. & Arlotta, P. 2014. Gene co-regulation by Fezf2 selects neurotransmitter identity and connectivity of corticospinal neurons. Nat Neurosci, 17, 1046–54.

Magavi, S. S. & Macklis, J. D. 2008. Identification of newborn cells by BrdU labeling and immunocytochemistry in vivo. Methods Mol Biol, 438, 335–43.

Mckenna, W. L., Betancourt, J., Larkin, K. A., Abrams, B., Guo, C., Rubenstein, J. L. & Chen, B. 2011. Tbr1 and Fezf2 regulate alternate corticofugal neuronal identities during neocortical development. J Neurosci, 31, 549–64.

Molyneaux, B. J., Arlotta, P., Hirata, T., Hibi, M. & Macklis, J. D. 2005. Fezl is required for the birth and specification of corticospinal motor neurons. Neuron, 47, 817–31.

Molyneaux, B. J., Goff, L. A., Brettler, A. C., Chen, H. H., Hrvatin, S., Rinn, J. L. & Arlotta, P. 2015. DeCoN: genome-wide analysis of in vivo transcriptional dynamics during pyramidal neuron fate selection in neocortex. Neuron, 85, 275–288.

Paxinos, G., Franklin, K.B.J 2001. The Mouse Brain in Stereotaxic Coordinates Elsevier.

Ptashne, M. 2014. The chemistry of regulation of genes and other things. J Biol Chem, 289, 5417–35.

Rouaux, C. & Arlotta, P. 2013. Direct lineage reprogramming of post-mitotic callosal neurons into corticofugal neurons in vivo. Nat Cell Biol, 15, 214–21.

Rouaux, C., Bhai, S. & Arlotta, P. 2012. Programming and reprogramming neuronal subtypes in the central nervous system. Dev Neurobiol, 72, 1085–98.

Shen, Q., Wang, Y., Dimos, J. T., Fasano, C. A., Phoenix, T. N., Lemischka, I. R., Ivanova, N. B., Stifani, S., Morrisey, E. E. & Temple, S. 2006. The timing of cortical neurogenesis is encoded within lineages of individual progenitor cells. Nat Neurosci, 9, 743–51.

Shimizu, T. & Hibi, M. 2009. Formation and patterning of the forebrain and olfactory system by zinc-finger genes Fezf1 and Fezf2. Dev Growth Differ, 51, 221–31.

Sugimoto, M., Fujikawa, A., Womack, J. E. & Sugimoto, Y. 2006. Evidence that bovine forebrain embryonic zinc finger-like gene influences immune response associated with mastitis resistance. Proc Natl Acad Sci U S A, 103, 6454–9.

Tasic, B., Menon, V., Nguyen, T. N., Kim, T. K., Jarsky, T., Yao, Z., Levi, B., Gray, L. T., Sorensen, S. A., Dolbeare, T., Bertagnolli, D., Goldy, J., Shapovalova, N., Parry, S., Lee, C., Smith, K., Bernard, A., Madisen, L., Sunkin, S. M., Hawrylycz, M., Koch, C. & Zeng, H. 2016. Adult mouse cortical cell taxonomy revealed by single cell transcriptomics. Nat Neurosci, 19, 335–46.

Tomassy, G. S., De Leonibus, E., Jabaudon, D., Lodato, S., Alfano, C., Mele, A., Macklis, J. D. & Studer, M. 2010. Area-specific temporal control of corticospinal motor neuron differentiation by COUP-TFI. Proc Natl Acad Sci U S A, 107, 3576–81.

Tsyporin, J., Tastad, D., Ma, X., Nehme, A., Finn, T., Huebner, L., Liu, G., Gallardo, D., Makhamreh, A., Roberts, J. M., Katzman, S., Sestan, N., Mcconnell, S. K., Yang, Z., Qiu, S. & Chen, B. 2021. Transcriptional repression by FEZF2 restricts alternative identities of cortical projection neurons. Cell Rep, 35, 109269.

Wenick, A. S. & Hobert, O. 2004. Genomic cis-regulatory architecture and trans-acting regulators of a single interneuron-specific gene battery in C. elegans. Dev Cell, 6, 757–70.

Wheat, J. C., Krause, D. S., Shin, T. H., Chen, X., Wang, J., Ding, D., Yamin, R. & Sweetser, D. A. 2014. The corepressor Tle4 is a novel regulator of murine hematopoiesis and bone development. PLoS One, 9, e105557.

Woodworth, M. B., Greig, L. C., Liu, K. X., Ippolito, G. C., Tucker, H. O. & Macklis, J. D. 2016. Ctip1 Regulates the Balance between Specification of Distinct Projection Neuron Subtypes in Deep Cortical Layers. Cell Rep, 15, 999–1012.

Xing, S., Shao, P., Li, F., Zhao, X., Seo, W., Wheat, J. C., Ramasamy, S., Wang, J., Li, X., Peng, W., Yu, S., Liu, C., Taniuchi, I., Sweetser, D. A. & Xue, H. H. 2018. Tle corepressors are differentially partitioned to instruct CD8(+) T cell lineage choice and identity. J Exp Med, 215, 2211–2226.

Yao, J., Liu, Y., Husain, J., Lo, R., Palaparti, A., Henderson, J. & Stifani, S. 1998. Combinatorial expression patterns of individual TLE proteins during cell determination and differentiation suggest non-redundant functions for mammalian homologs of Drosophila Groucho. Dev Growth Differ, 40, 133–46.

Zhang, S., Li, J., Lea, R., Vleminckx, K. & Amaya, E. 2014. Fezf2 promotes neuronal differentiation through localised activation of Wnt/β-catenin signalling during forebrain development. Development, 141, 4794–805.

Ziller, M. J., Edri, R., Yaffe, Y., Donaghey, J., Pop, R., Mallard, W., Issner, R., Gifford, C. A., Goren, A., Xing, J., Gu, H., Cachiarelli, D., Tsankov, A., Epstein, C., Rinn, J. R., Mikkelsen, T. S., Kohlbacher, O., Gnirke, A., Bernstein, B. E., Elkabetz, Y. & Meissner, A. 2015. Dissecting neural differentiation regulatory networks through epigenetic footprinting. Nature, 518, 355–359.

