## Supplemental Material_Galazo, Sweetser, Macklis for "*Tle4* controls both developmental acquisition and postnatal maintenance of corticothalamic projection neuron identity"

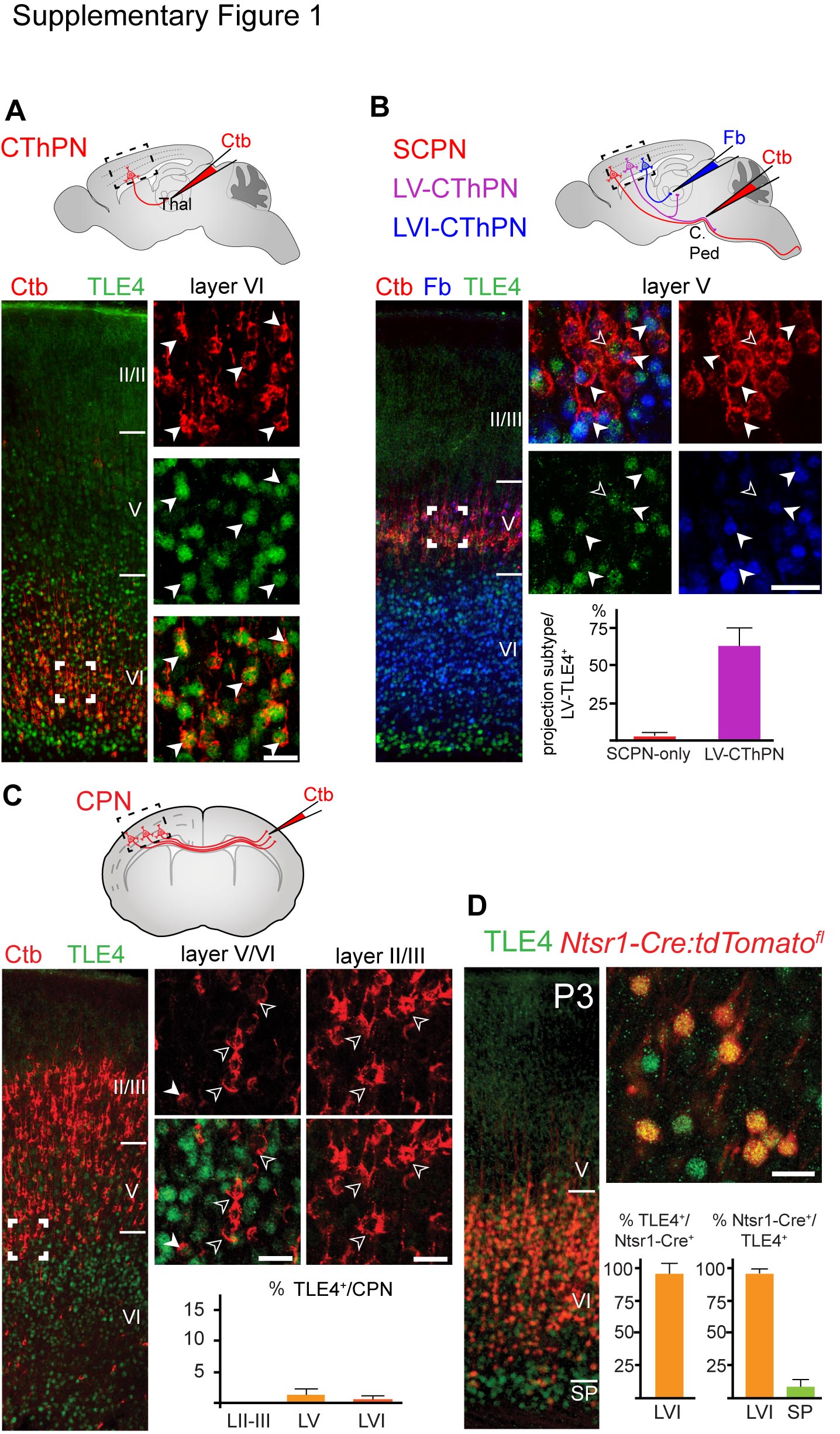

**Supplementary Figure 1**. ***Tle4* is constitutively expressed by Layer VI-CThPN and Layer V-CThPN and excluded from other subtypes**

TLE4 is expressed by both functional subtypes of corticothalamic neurons– layer VI CThPN that project only to the thalamus, and layer V CThPN that project to both thalamus and subcrebrally– but it is only rarely expressed by other projection neuron subtypes. Schematics depict retrograde labeling approaches for neuron subtype identification at the top of each panel.

(A) TLE4 immunocytochemistry (green) on coronal sections from P6 brains injected with cholera toxin-b (Ctb; red) in the thalamus. All TLE4^+^ neurons in layer VI (green) are labeled with retrograde Ctb (red). (B) Double retrograde labeling approach to identify dual projecting layer V-CThPN. Fast blue (blue) was injected into the thalamus, and Ctb (red) was injected into the cerebral peduncle. TLE4 co-localization with Fb and Ctb reveals that 62.2% of TLE4^+^ neurons in layer V are dual projecting CThPN (double-labeled Fb^+^-Ctb^+^; white arrowheads). Only 2.5% of TLE4^+^ neurons in layer V are Ctb^+^-only (SCPN, open arrowheads). (C) Almost no callosal projection neurons (CPN; Ctb^+^) express TLE4 (open arrowheads). Only rare TLE4^+^ CPN (Ctb^+^) are present in deep layers (solid arrow) with none in superficial layers. (D) TLE4 immunolabeling in *Ntsr1-Cre:tdTomato^fl^* cortex at P3. All Cre-expressing neurons (tdTomato^+^) are located in layer VI and subplate, but not in layer V. Virtually all *Cre*-expressing neurons are TLE4^+^ (96%), and 94% of TLE4+ neurons in layer VI express *Cre*, but only 6% of TLE4^+^ in subplate neurons express *Cre*. Scale bars, 20 μm (A-D).

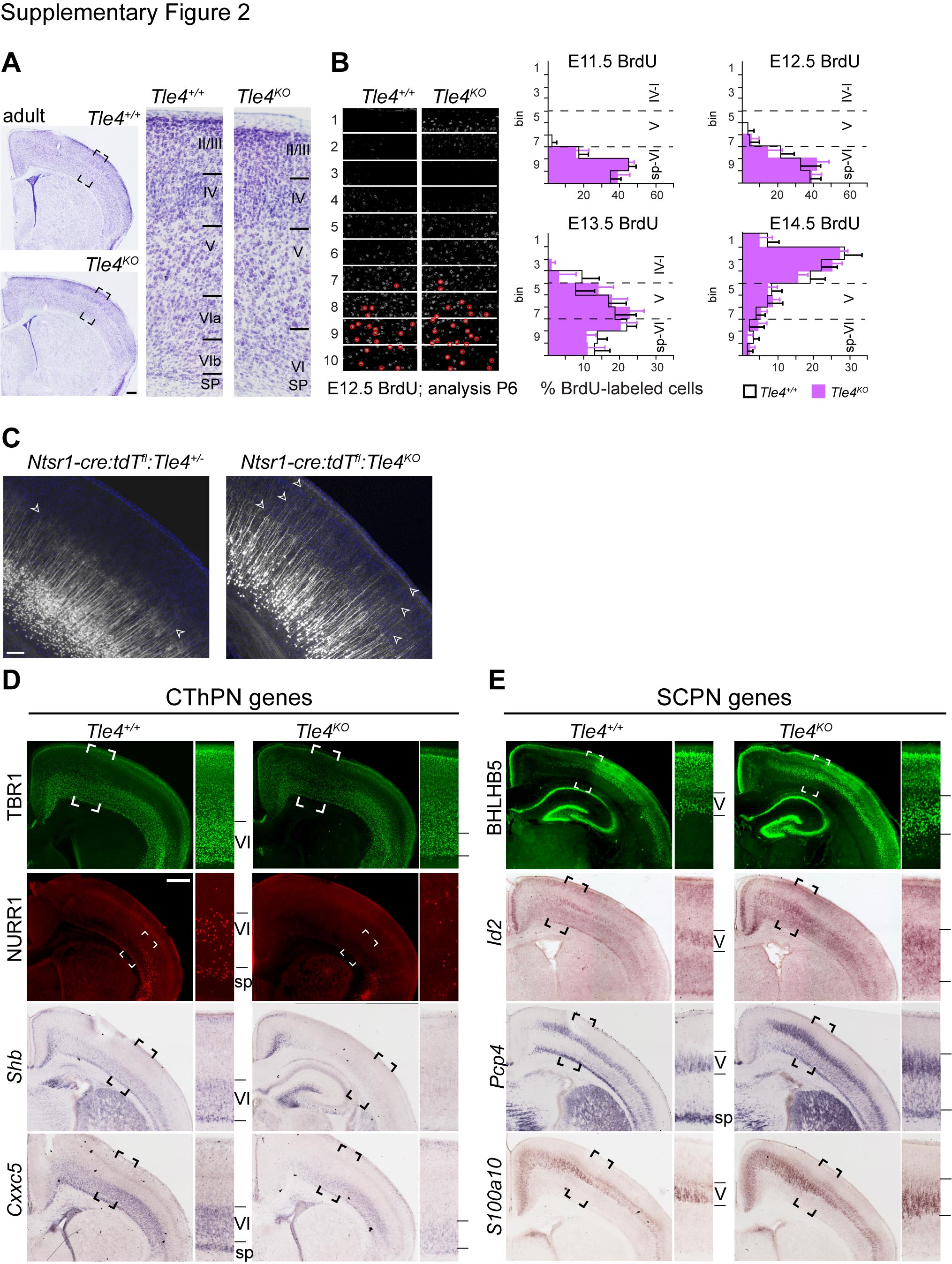
**Supplementary Figure 2. Cytoarchitectural changes in *Tle4^KO^* cortex are not due to abnormal generation or distribution of early born neurons, but instead reflect abnormal acquisition of CThPN molecular identity and dendritic connectivity**

(**A**) Expansion of layer V and contraction of layer VI persist in adult *Tle4^KO^* cortex. Nissl straining shows cortical cytoarchitecture of 3 month-old wildtype and *Tle4^KO^* mice. (**B**) The number and laminar distribution at P6 of cells labeled by BrdU at E11.5, E12.5, E13.5, and E14.5 is similar in *Tle4^KO^* and wildtype mice. Bins 1-4 correspond to layers I-IV, bins 5-7 correspond to layer V, and bins 8-10 correspond to layer VI and subplate. (**C**) More neurons extend apical dendrites into superficial layers II/III and layer I in *Tle4^KO^* mice compared to wildtypes. Dendrites are visualized by tdTomato endogenous fluorescence from the CThPN-specific reporter *Ntsr1-Cre:tdTomato^fl^.* Open arrowheads indicate cortical depths reached by CThPN apical dendrites at P6. (D-E) In the absence of *Tle4* function, expression of CThPN genes TBR1, NURR1, *Shb*, and *Cxxc5* decreases (D), and expression of SCPN genes BHLHB5, *Id2, Pcp4,* and *S100a10* increases at P8 (E). Insets correspond to the boxed areas in the low magnification images. Scale bars, 500 μm (A), 100 μm (C), 500 μm (D-E).

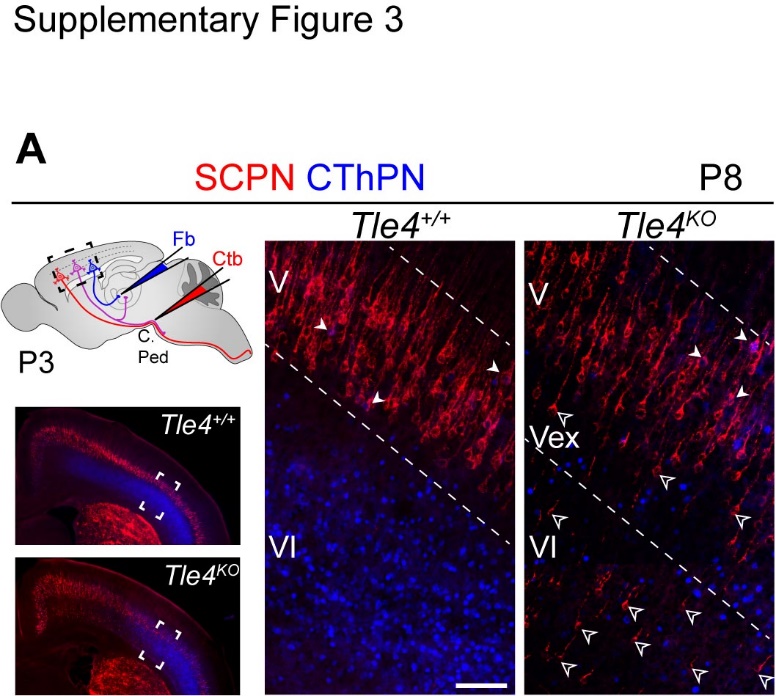

**Supplementary Figure 3.** **In *Tle4^KO^* cortex, projections to the thalamus are reduced, since fate-converted SCPN do not develop a thalamic projection.**

(A) Double retrograde labeling from the thalamus and cerebral peduncle at P3 to identify neurons with dual projections in control and *Tle4^KO^* mice. Fb (blue) was injected into the thalamus, and Ctb (red) into the cerebral peduncle. In *Tle4^KO^* and control mice, neurons with dual projections (Fb^+^-Ctb^+^) are located only in layer V, the normal location of dual-projecting layer V-CThPN (white arrowheads). No fate-converted SCPN (Ctb^+^ in layers V_ex_ or VI) in Tle4^KO^ mice are double labeled (open arrowheads). High magnification images correspond to the boxed areas. Scale bar, 100 μm.

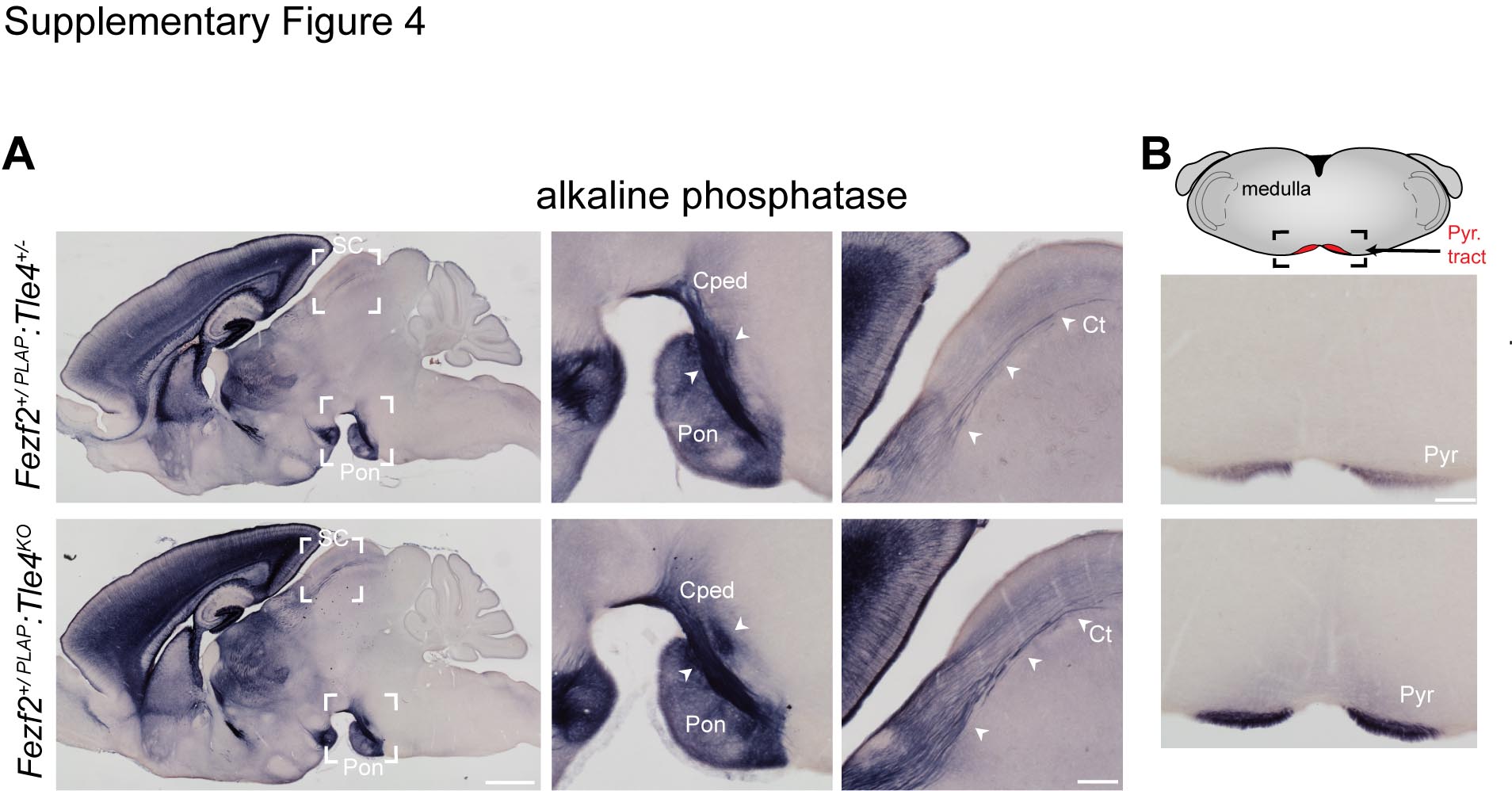

**Supplementary Figure 4. Increased projection to subcerebral targets, but no frankly ectopic axon projections are present in** ***Tle4^KO^* mice**

(**A**) Visualization of axonal projections from all *Fezf2*-expressing neurons at P21 using *Fezf2*^+^*^/PLAP^* as the reporter. Insets show the boxed areas at higher magnification, corresponding to the cerebral peduncle (Cped) and pontine nuclei (Pon), and the corticotectal tract (Ct) projecting into the superior colliculus (SC). More PLAP-labeled axons are present in the cerebral peduncle and corticotectal tracts in *Fezf2*^+^*^/PLAP^:Tle4*^KO^ mice compared to *Fezf2*^+^*^/PLAP^:Tle4*^+/-^ mice. White arrowheads mark PLAP-labeled projections in these areas. (**B**) Increased PLAP labeling in the pyramidal tract (Pyr) reveals increased subcerebral projection at the level of medulla, *en route* to the spinal cord. Scale bar, 1mm (A), 250 μm (A, insets), 250 μm (B).

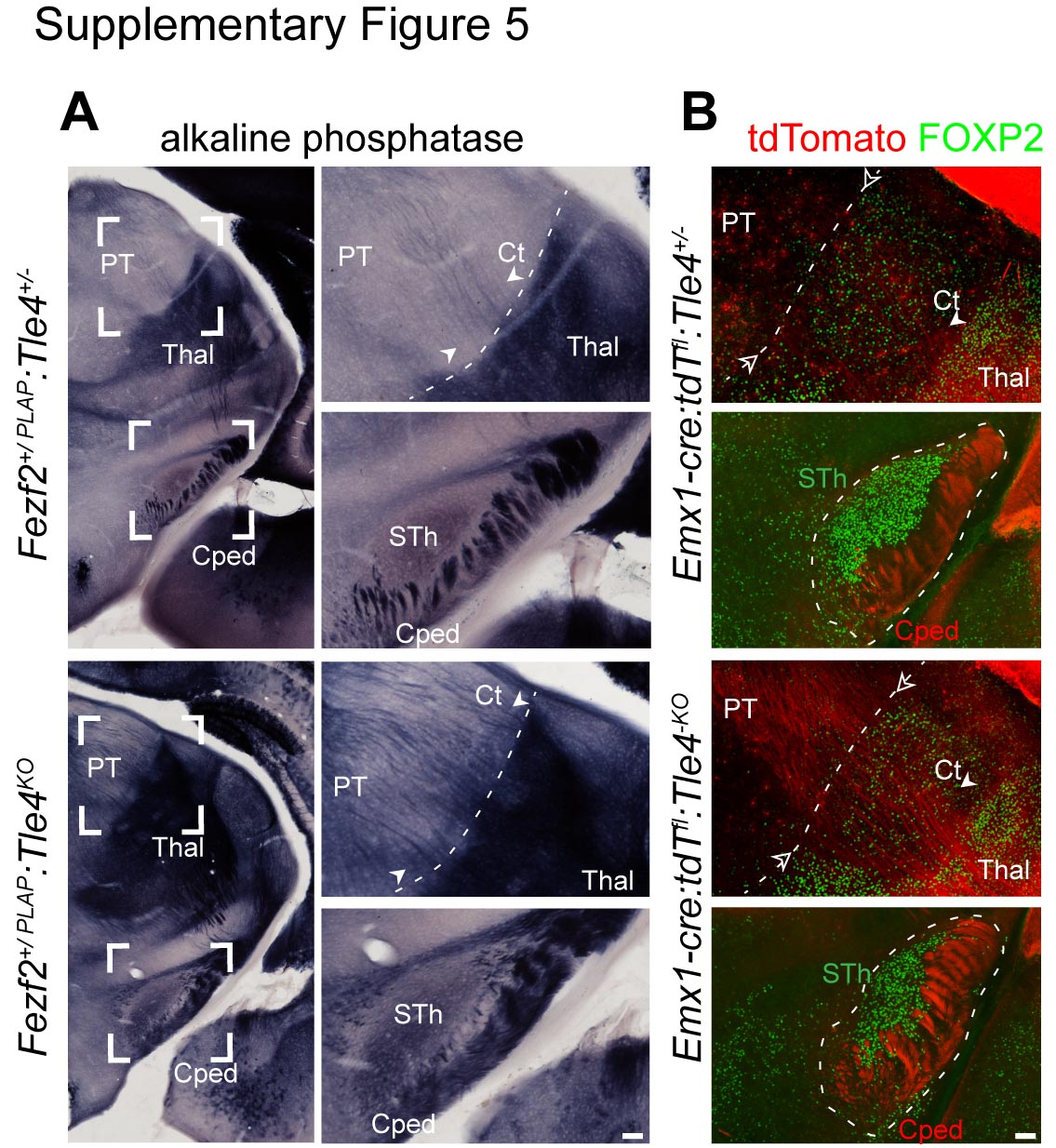

**Supplementary Figure 5**. **Increased projection to the cerebral peduncle and superior colliculus in *Tle4^KO^* mice compared to controls**

(**A**) Visualization of axonal projections from *Fezf2*-expressing neurons at P21 using a *Fezf2*^+^*^/PLAP^* reporter. Insets show at higher magnification the boxed areas of the cerebral peduncle (Cped), at the level of the subthalamic nucleus (STh), and of the border between thalamus (Thal) and pretectal area (PT) (dashed line). Corticotectal axons (Ct) crossing the thalamic border *en route* to the superior colliculus are indicated by white arrowheads. PLAP labeling reveals that more axons project into PT *en route* to the superior colliculus, and into the cerebral peduncle in *Fezf2*^+^*^/PLAP^:Tle4*^KO^ mice compared to *Fezf2*^+^*^/PLAP^:Tle4*^+/-^ mice. (**B**) Visualization of subcerebral projections using cortex-specific reporter *Emx1-Cre:tdTomato^fl^* at P14 in areas equivalent to those shown in (A) with *Fezf2*^+^*^/PLAP^*. Increased projections en route to the superior colliculus and into the cerebral peduncle revealed by tdTomato fluorescence in *Emx1-Cre:tdTomato^fl^:Tle4^KO^* mice compared to *Emx1-Cre:tdTomato^fl^:Tle4^+/-^* mice. FOXP2 immunolabeling (green) is used to delineate the border between thalamus and PT (dashed line), and the subthalamic nucleus (STh). Scale bars, 100 μm (A-B).

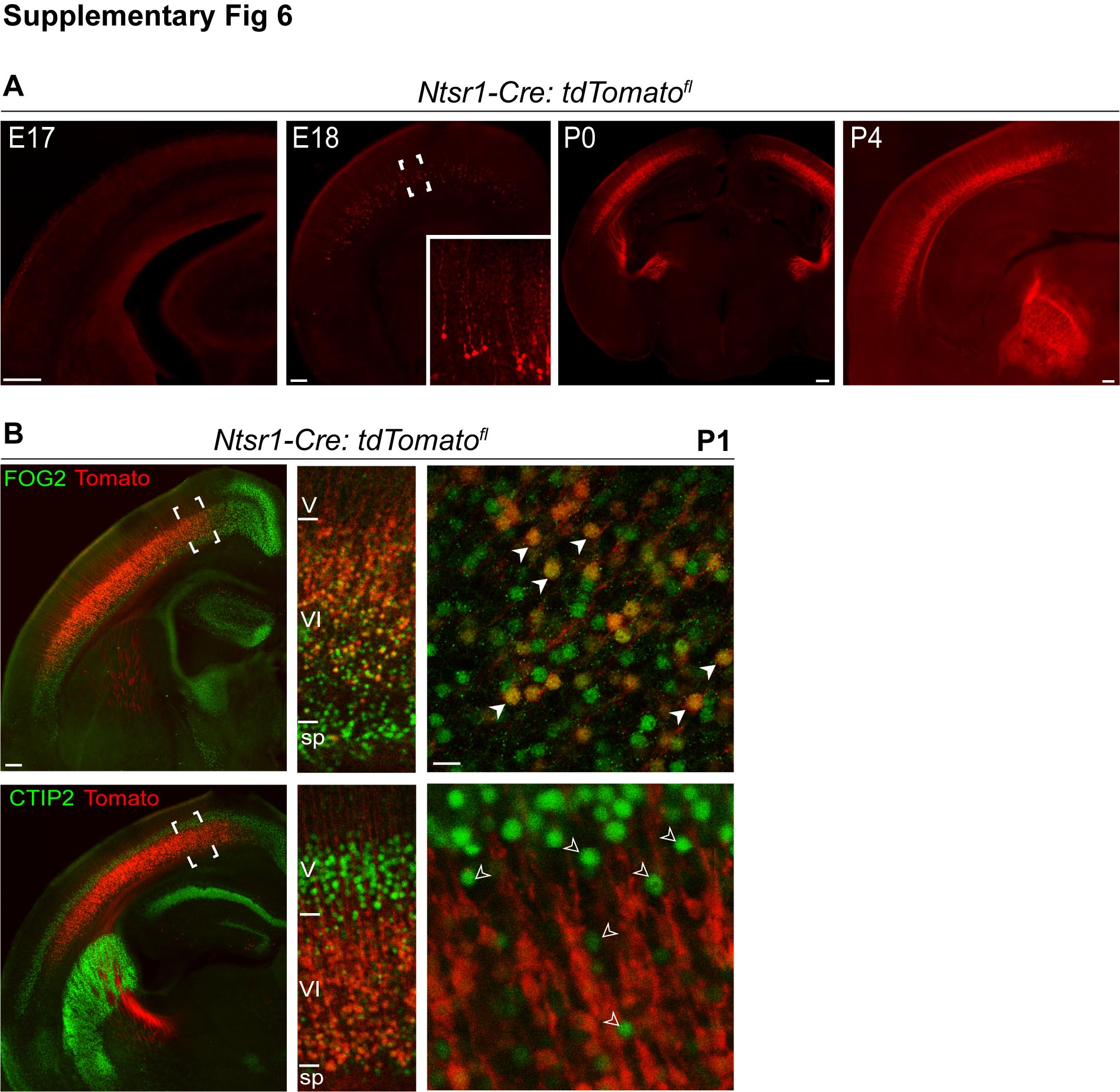

**Supplementary Figure 6.** **Time course of *Cre* expression in Ntsr1-Cre mice, and specificity for CThPN perinatally**

(**A**) Developmental dynamics of *Cre* expression revealed by tdTomato reporter in *Ntsr1-Cre:tdTomato^fl^* mice. Before E18, expression is negligible. At E18, *Cre* expression is detected in neurons with pyramidal morphology and apical dendrites in developing layer VI. Inset shows higher magnification of boxed area. At P0, layer VI CThPN have robust *Cre* expression. TdTomato reveals CThPN projections entering the thalamus at P0, and more developed projections at P4. (**B**) Immunolabeling for FOG2 and CTIP2 in *Ntsr1-Cre:tdTomato^fl^* cortices at P1. TdTomato reporter fluorescence co-localizes with FOG2 (white arrowheads), but not with CTIP2 (open arrowheads), indicating that *Cre* is expressed specifically by CThPN, and is not expressed by SCPN. Scale bars, 200 μm (A), 200 μm (B, mid magnification), 20 μm (B, high magnification).

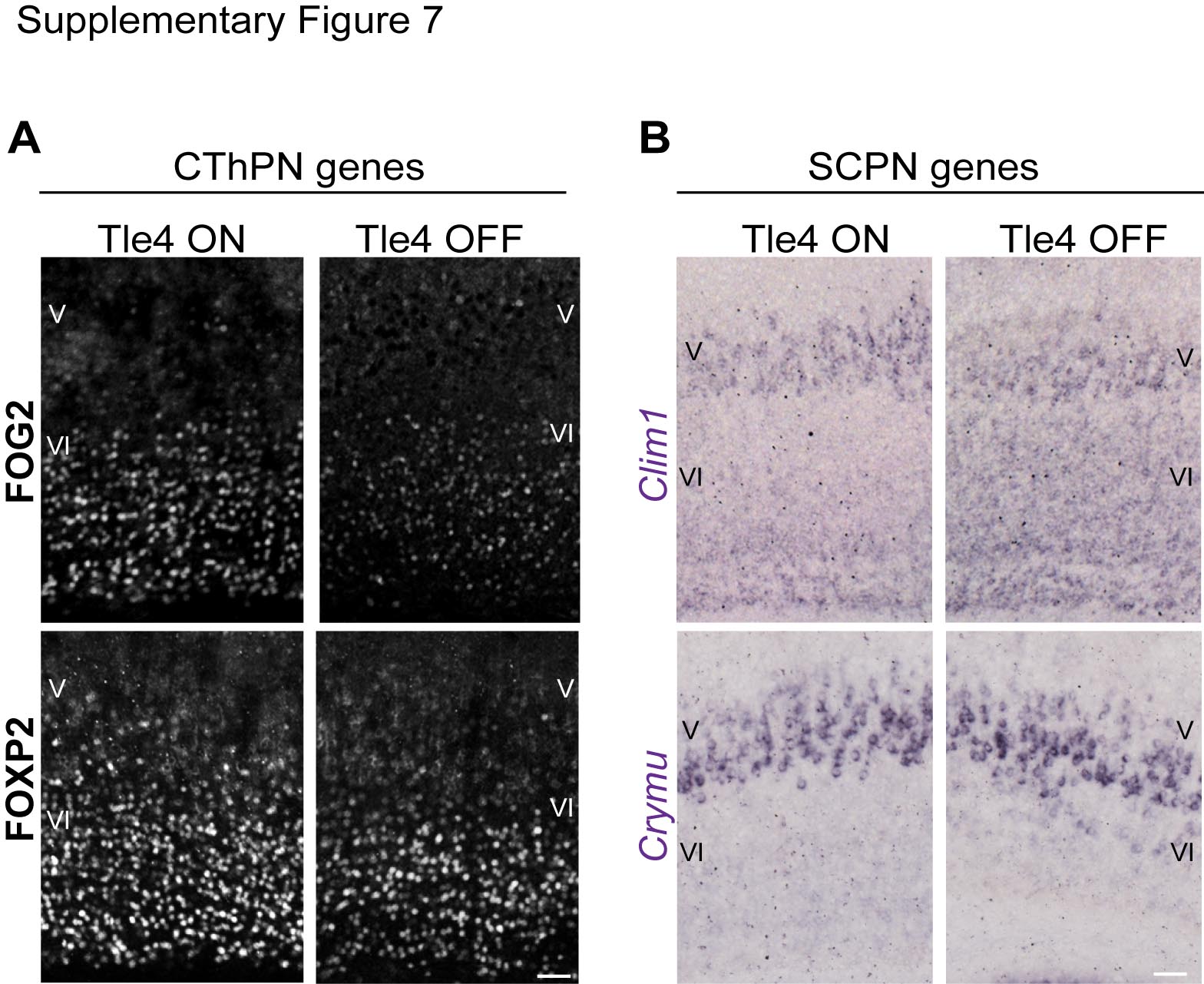

**Supplementary Figure 7.** **Loss of** ***Tle4* function during CThPN maturation results in downregulation of cardinal CThPN genes and upregulation of cardinal SCPN genes by differentiated CThPN**

(**A**) Immunolabeling for FOG2 and FOXP2 shows downregulation of CThPN genes in layer VI of a cortical area in which *Tle4* expression has been silenced after AAV-Cre injection at P3 (Tle4 OFF). In the contralateral control hemisphere (Tle4 ON), *Tle4* expression is normal. (**B**) ISH reveals strong upregulation of *Clim1*, and modest upregulation of *Crymu*, in layer VI of the cortical area injected with AAV-Cre at P3, but not in the contralateral control hemisphere. Scale bars, 50 μm (A, B).

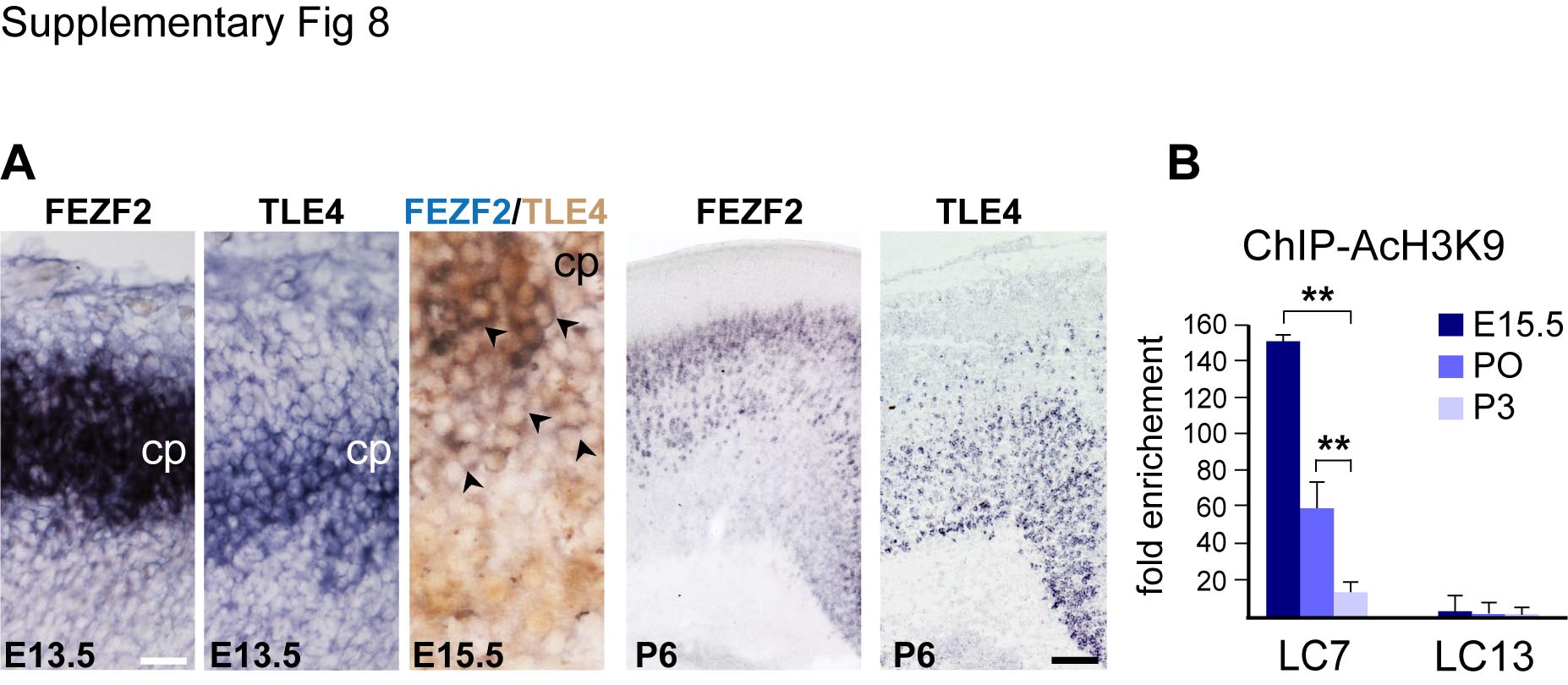
**Supplementary Figure 8. Early born cortical neurons express *Fezf2* and *Tle4* strongly during embryonic development, but *Fezf2* expression by CThPN substantially diminishes during postnatal maturation**

(**A**) ISH for *Fezf2* and *Tle4* in adjacent 10 μm sections reveals that both are expressed strongly by postmitotic neurons in the cortical plate at E13.5. Combined *Fezf2* ISH with TLE4 immunolabeling directly demonstrates co-localization of *Fezf2* and TLE4 in cortical plate neurons (black arrowheads) at E15.5. At P6, *Fezf2* and *Tle4* are most strongly expressed in layer V and layer VI, respectively. (**B**) ChIP-AcH3K9 reveals that the acetylation level of Locus 7, co-occupied by TLE4 and FEZF2, progressively decreases during embryonic and postnatal development, while the acetylation level of Locus 13, not co-occupied by TLE4 and FEZF2, is extremely low and does not change over time. ChIP-AcH3K9 fold enrichment was determined by normalizing ΔCt values for each locus against the ΔCt values for a constitutively active locus (*Gapdh*). Asterisks indicate significance (two-tailed t-test with Tukey correction for pairwise comparisons; **p<0.01). Scale bars, 20 μm (A, high magnification) and 100 μm (A, low magnification).

**Supplementary Table 1. ChIP-qPCR loci coordinates**

Coordinates in chromosome 14 corresponding to the loci amplified for ChIP-FEZF2 and ChIP-TLE4 experiments. UCSC genome browser was used to map coordinates to the GRCm38/mm10 genome assembly.

| **ChIP-qPCR**  **Loci** | **Chromosome 14 coordinates** |
| --- | --- |
| ***LC1*** | 12342133..12324458 |
| ***LC2*** | 12343008..12342368 |
| ***LC3*** | 12341906..12342164 |
| ***LC4*** | 12341756..12341031 |
| ***LC5*** | 12341516..12341813 |
| ***LC6*** | 12341219..12341454 |
| ***LC7*** | 12341036..12341348 |
| ***LC8*** | 12340849..12341220 |
| ***LC9*** | 12340723..12341013 |
| ***LC10*** | 12340435..12340793 |
| ***LC11*** | 12340147..12340521 |
| ***LC12*** | 12340000..12340302 |
| ***LC13*** | 12339795..12340164 |
| ***LC14*** | 12339553..12339857 |
| ***LC15*** | 12339280..12339606 |
| ***LC16*** | 12339021..12339386 |
| ***LC17*** | 12338844..12339231 |
| ***LC18*** | 12338683..12339037 |
| ***LC19*** | 12338361..12338648 |
| ***LC20*** | 12338200..12338505 |
| ***LC21*** | 12338100..12338381 |
| ***LC22*** | 12337734..12338100 |
| ***LC23*** | 12337567..12337900 |
